# Massively parallel, time-resolved single-cell RNA sequencing with scNT-Seq

**DOI:** 10.1101/2019.12.19.882050

**Authors:** Qi Qiu, Peng Hu, Kiya W. Govek, Pablo G. Camara, Hao Wu

## Abstract

Single-cell RNA sequencing offers snapshots of whole transcriptomes but obscures the temporal dynamics of RNA biogenesis and decay. Here we present single-cell new transcript tagging sequencing (scNT-Seq), a method for massively parallel analysis of newly-transcribed and pre-existing RNAs from the same cell. This droplet microfluidics-based method enables high-throughput chemical conversion on barcoded beads, efficiently marking metabolically labeled newly-transcribed RNAs with T-to-C substitutions. By simultaneously measuring new and old transcriptomes, scNT-Seq reveals neuronal subtype-specific gene regulatory networks and time-resolved RNA trajectories in response to brief (minutes) versus sustained (hours) neuronal activation. Integrating scNT-Seq with genetic perturbation reveals that DNA methylcytosine dioxygenases may inhibit stepwise transition from pluripotent embryonic stem cell state to intermediate and totipotent two-cell-embryo-like (2C-like) states by promoting global RNA biogenesis. Furthermore, pulse-chase scNT-Seq enables transcriptome-wide measurements of RNA stability in rare 2C-like cells. Time-resolved single-cell transcriptomic analysis thus opens new lines of inquiry regarding cell-type-specific RNA regulatory mechanisms.

## INTRODUCTION

Dynamic changes in RNA levels are regulated by the interplay of RNA transcription, processing, and degradation^1, 2^. Understanding the mechanisms by which the transcriptome is regulated in functionally diverse cell-types within multi-cellular organisms thus requires cell-type-specific measurements of the kinetics of RNA biogenesis and decay. Recent advances in single-cell RNA sequencing (scRNA-Seq) technologies are leading to a more complete understanding of heterogeneity in cell types and states^3^. However, standard scRNA-Seq methods capture a mixture of newly-synthesized (“new”) and pre-existing (“old”) RNAs without being able to temporally resolve RNA dynamics.

Accurately capturing changes in the nascent transcriptome over time at single-cell resolution is of particular interest because measuring newly-transcribed RNAs can reveal immediate regulatory changes in response to developmental, environmental, metabolic, and pathological signals^4^. Commonly used approaches for distinguishing newly-transcribed from pre-existing RNAs of the same population of transcripts rely on metabolic labeling that employs thiol-labeled nucleoside analogs such as 4-thiouridine (4sU) and subsequent biochemical enrichment of metabolically labeled RNAs^2^. Although these methods have yielded unprecedented insights into the regulation of RNA dynamics, they require ample starting material and present challenges for enrichment normalization. Several approaches have recently been developed to chemically convert 4sU into cytidine analogs, yielding uracil-to-cytosine (U-to-C) substitutions that label newly-transcribed RNAs after reverse transcription^5–7^. These chemical nucleotide conversion methods allow for direct measurement of temporal information about cellular RNAs in a sequencing experiment without biochemical enrichment. Recent studies integrated standard plate-based single-cell RNA-Seq method with one of these chemical methods (i.e. thiol(SH)-linked alkylation for the metabolic sequencing of RNA (SLAM)-Seq)^8, 9^, demonstrating the feasibility of studying newly-transcribed transcriptomes at single-cell levels. However, these plate-based single-cell SLAM-Seq methods suffer from several limitations. First, they are costly, and the associated library preparation steps are time-consuming, prohibiting it for large-scale single-cell analysis of highly heterogeneous cell populations. Second, these methods lack unique molecular identifiers (UMIs), preventing accurate quantification of the newly-transcribed transcript levels.

To overcome these constraints, we developed single-cell new transcript tagging sequencing (scNT-Seq), a high-throughput and UMI-based scRNA-Seq method that simultaneously measures both new and old transcriptomes from the same cell. In scNT-Seq, integration of metabolic RNA labeling, droplet microfluidics, and chemically induced recoding of 4sU to cytosine analog permits highly scalable and time-resolved single-cell analysis of cellular RNA dynamics. We demonstrate that the method is easy to set up and substantially improves the time and cost. We show scNT-Seq enables more detailed characterization of gene regulatory networks and temporal RNA trajectories than single-cell whole-transcriptome measurements alone.

## RESULTS

### Development and validation of scNT-Seq

To enable newly-transcribed and pre-existing transcripts from the same cell to be analyzed in a scalable manner, we focused on the Drop-Seq platform because its unique barcoded bead design affords both immobilization of RNAs for efficient chemical conversion reactions and UMI-based high-throughput scRNA-Seq analysis, and this droplet microfluidics platform has been widely adopted^10–14^. The scNT-Seq consists of the following key steps (**Fig. 1a**): (1) metabolically labeling of cells with 4sU for a temporally defined time period; (2-3) co-encapsulating each individual cell with a barcoded oligo-dT primer coated bead in a nanoliter-scale droplet which captures both newly-transcribed and pre-existing RNAs; (4) performing 4sU chemical conversion on pooled barcoded beads in one reaction after droplet breakage; (5-8) reverse transcription, cDNA amplification, tagmentation, indexing PCR, and sequencing; and (9) using a UMI-based statistical model to analyze T-to-C substitutions within transcripts and infer the new transcript fraction at single-cell level.

**Figure 1.**
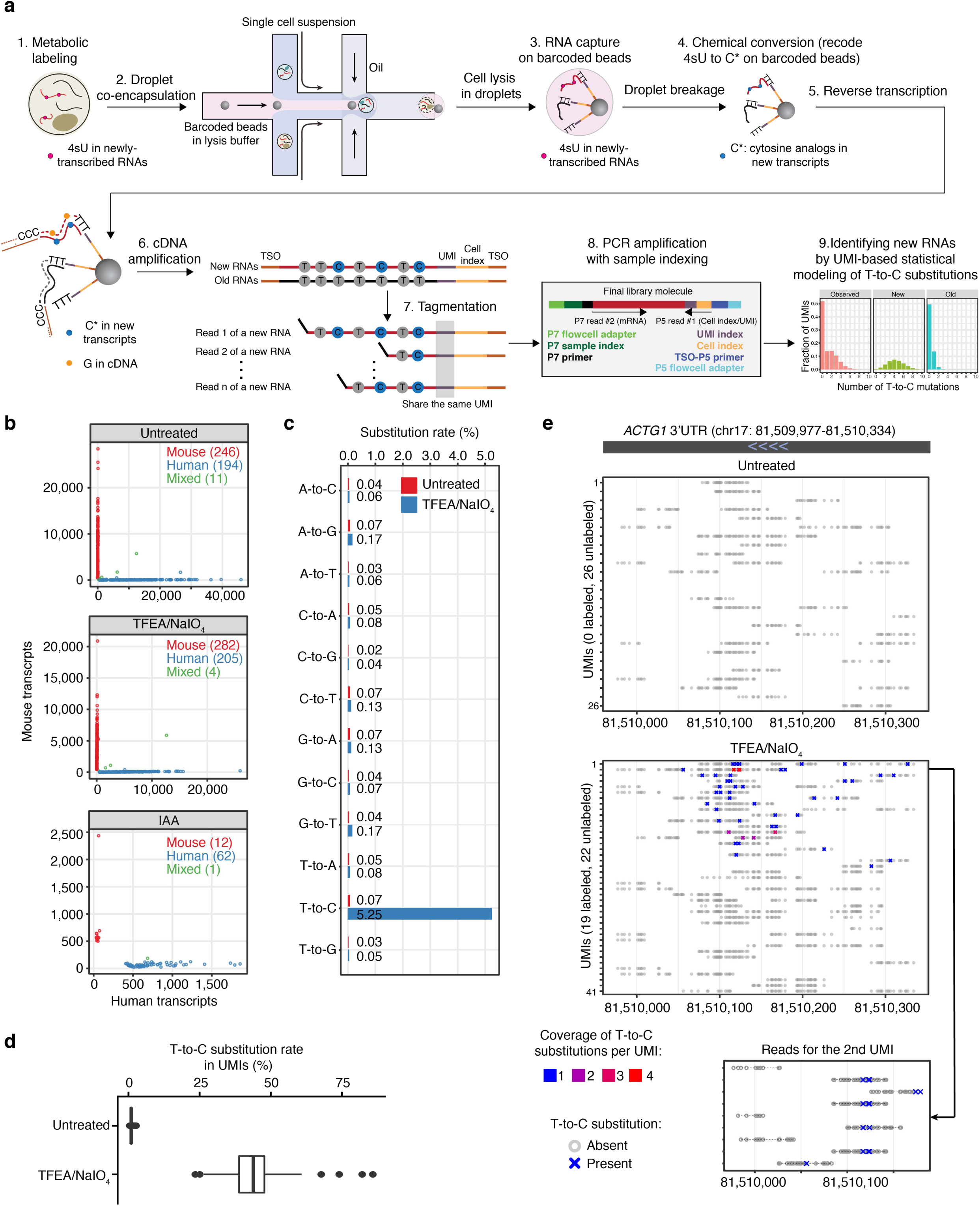
Development and validation of scNT-Seq. (**a**) Overview of <u>s</u>ingle-<u>c</u>ell <u>N</u>ascent transcript <u>T</u>agging sequencing (scNT-Seq). (**b**) Multi-species mixing experiment measures scNT-Seq specificity. Mouse ESCs and human K562 cells were mixed at 1:1 ratio after 4sU labeling (100 μM, 4 hrs). Scatterplot shows the number of transcripts (UMIs) mapped to mouse (y-axis) or human (x-axis) genome for each cell (dot). Cells with mostly mouse transcripts are labeled as mouse (red), while cells with mostly human transcripts are labeled as human (blue). Cells with a relatively high percentage of both mouse and human transcripts are labeled as mixed (green). (**c**) Transcriptome-wide nucleotide substitutions rates demonstrate specific T-to-C conversion in 4sU labeled K562 cells after TEFA/NaIO4-treatment compared to untreated control. (**d**) Box plots showing percentage of T-to-C containing transcripts (UMIs) in individual K562 cells. The box plots display the median (center line) and interquartile range (IQR, from the 25th to 75th percentile), the whiskers represent 1.5 times the interquartile range, and the circles represent outliers. (**e**) All UMIs of the *ACTG1* gene are shown for one K562 cell with or one without TFEA/NaIO4-treatment. Grey circles stand for T without T-to-C substitution, while crosses (“X”s) stand for sites of T-to-C substitution in at least one read. The color of “X” indicates the read coverage in a UMI containing T-to-C substitution at a particular site. All reads from one exemplary transcript (2^nd^ UMI from TFEA/NaIO4-treated sample) are further shown in the lower panel.

To identify the optimal reaction conditions on barcoded beads, we first explored two independent chemical conversion methods (TimeLapse-Seq: 2,2,2-trifluoroethylamine (TFEA)/sodium periodate (NaIO4)-based 4sU conversion; SLAM-Seq: iodoacetamide (IAA)-based conversion) and validated their performance with species-mixing experiments using cultured mouse embryonic stem cells (mESCs) and human K562 cells. This analysis indicates that TFEA/NaIO4-based scNT-Seq substantially outperforms the IAA-based assay in terms of mRNA recovery rates (**Fig. 1b**). Furthermore, the collision rate is comparable between TFEA/NaIO4-based scNT-Seq and standard Drop-Seq (**Fig. 1b**), demonstrating the specificity of scNT-Seq in analyzing single-cell transcriptomes. Shallow sequencing of mESCs under different treatment conditions showed that TFEA/NaIO4-based scNT-Seq identified a similar number of genes or UMIs per cell compared to standard Drop-Seq (**Supplementary Fig. 1a**). Aggregated single-cell new or old transcriptomes were highly correlated between replicates (**Supplementary Fig. 1b**). Further analysis of K562 cells revealed that only 4sU labeling and TFEA/NaIO4 treatment resulted in a substantial increase in T-to-C substitution (**Fig. 1c**) and in UMIs that contained one or more such substitutions (**Fig. 1d, e**). Notably, TFEA/NaIO4 chemical treatment works efficiently with both freshly isolated and cryo-preserved (methanol-fixed) cells (**Supplementary Fig. 1c**), demonstrating the versatility of scNT-Seq. Collectively, these data indicate that TFEA/NaIO4-based scNT-Seq is capable of efficiently detecting newly synthesized transcripts at single-cell resolution.

### Application of scNT-Seq to study neuronal activity-dependent transcription and regulatory networks

Neuronal activity induces expression of hundreds of activity-regulated genes (ARGs) in the vertebrate brain, leading to new protein synthesis and epigenetic changes necessary for short- and long-term memories of experiences. Thus, the coupling of synaptic activity to nascent transcription in the nucleus allows neurons to both respond dynamically to their immediate environment, and to store information stably^15^. Recent studies suggest that different patterns of neuronal activity could induce a distinct set of ARGs^16^. To investigate cell-type-specific transcription in response to distinct activity-duration patterns (brief versus sustained stimulation), we applied scNT-Seq to primary cortical neuronal cultures, derived from mouse cortex (embryonic day 16 (E16)), which contains heterogeneous populations of neuronal subtypes and undifferentiated neural progenitor cells.

To detect activity-regulated immediate transcriptional responses, we metabolically labeled primary mouse cortical cultures (200 μM 4sU) for two hours and stimulated the cells with different durations of neuronal activity (0, 15, 30, 60 and 120 min of potassium chloride (KCl)-mediated membrane depolarization) (**Fig. 2a**). After filtering low-quality cells and potential doublets, we retained 20,547 single-cell transcriptomes from five time-points (**Fig. 2b** and **Supplementary Table 1**). After sample integration, principal component analysis (PCA), graph-based clustering, and visualization by Uniform Manifold Approximation and Projection (UMAP), we identified nine major cell-types based on known marker genes: *Neurod6*+ cortical excitatory neurons (Ex, 68.5%), four *Gad1*+ inhibitory neuronal subtypes (Inh1-4, 13.9% in total), *Dlx1/Dlx2*+ inhibitory neuronal precursors (Inh-NP: 1.7%), two sub-populations of *Nes/Sox2*+ excitatory neuronal precursors (Ex-NP1/2: 10.4% in total), and *Nes/Aldh1l1*+ radial glia (RG: 5.5%) (**Fig. 2b** and **Supplementary Fig. 2b, c**).

**Figure 2.**
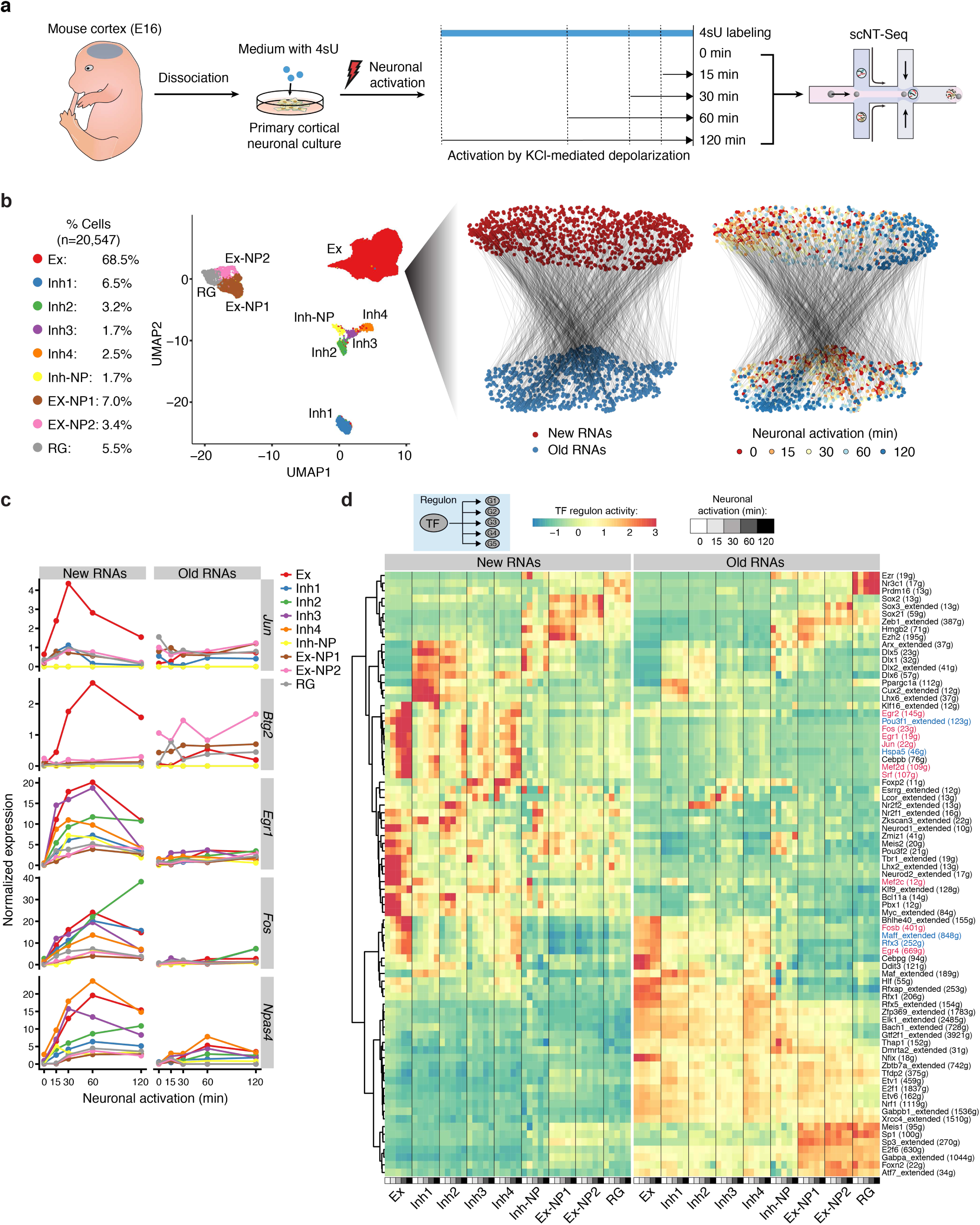
scNT-Seq captures cell-type-specific RNA dynamics and TF activity in response to distinct neuronal activity patterns. (**a**) Schematics of applying scNT-Seq analysis to study neuronal activation of primary mouse cortical cultures. (**b**) Uniform manifold approximation and projection (UMAP) visualization of 20,547 cells from primary mouse cortical cultures. Relative percentage of each cell-type is listed on the left. Randomly sampled 1,000 excitatory neurons from 5 time points (200 cells from each time point) are sub-clustered based on newly-synthesized (top) or pre-existing transcripts (bottom). Cells from each time point are color-coded and projected into the stacked UMAP on the right panel. The new and old transcriptomes of same cell were connected by a black line. Ex, excitatory neurons; Inh, inhibitory neurons; NP, neural progenitors; RG, radial glia. (**c**) Cell-type-specific responses of activity-regulated genes (ARGs) upon neuronal activation. The mean new and old RNA expression levels (transcripts per 10k, TP10k) were shown in line plot. (**d**) Heat map showing cell-type-specific TF regulon activity (AUC value quantified by SCENIC) in response to distinct activity durations. Red color highlights known regulators of ARGs, while blue color highlights potential novel regulators induced by neuronal activity.

We next sought to distinguish newly-transcribed from pre-existing RNAs by counting and statistically modeling T-to-C substitutions in transcripts (UMIs), an approach that overcomes the problem of incomplete 4sU labeling of new transcripts (up to 50% of all reads originated from new RNAs may not contain T-to-C substitutions)^17^. The quantification accuracy is further improved by UMI-based analysis, which substantially increases the number of uridines or T-to-C substitutions covered in each transcript compared to the analysis of individual sequencing reads (**Supplementary Fig. 3a, b**). After statistical correction, we obtained reliable measurements of newly-transcribed RNA faction for both activity-induced genes (e.g., *Fos*, ∼90% new/total) and slow turnover house-keeping genes (e.g., *Mapt*, <5% new/total) in excitatory neurons (**Supplementary Fig. 3c, d**).

Furthermore, sub-clustering of newly-transcribed (upper) or pre-existing (lower) transcriptomes derived from randomly sampled excitatory neurons could readily separate new, but not old single-cell transcriptomes in an activity-pattern dependent manner (right panels in **Fig. 2b**). In contrast, new transcriptomes of undifferentiated excitatory neural precursors (Ex-NP/RG) did not exhibit similar distributions (**Supplementary Fig. 4a**). Next, we directly examined how transcription of classic ARGs are induced in different cell-types in response to distinct activity-patterns. While some ARGs, such as *Jun* and *Btg2*, were specifically induced in Ex neurons upon activation, other ARGs (e.g., *Egr1*, *Fos*, and *Npas4*), were broadly induced in many cell-types including non-neuronal cells, albeit with different magnitudes and response curves (**Fig. 2c, Supplementary Fig. 4b**). There was little to no change at pre-existing levels upon activation of Ex neurons (**Fig. 2c**). Together, these results suggest scNT-Seq can accurately detect cell-type-specific, activity-induced immediate transcriptional changes within the timescale of minutes to hours.

Recent advances in computational analysis enable identification of specific gene regulatory networks (GRNs, also known as regulon of a transcription factor (TF)) underlying stable cell states by linking *cis*-regulatory sequences to single-cell gene expression. Since the regulon is scored as a whole, instead of using the expression of individual genes, this approach is more robust against experimental dropouts or stochastic variation of gene expression due to transcriptional bursting. We reasoned that applying such an approach to newly-transcribed RNAs derived from scNT-Seq may allow for analysis of regulons underlying dynamic cell states induced by external stimuli such as neuronal activity. By applying single-cell regulatory network inference and clustering (SCENIC)^18^ to statistically-corrected newly-transcribed and pre-existing single-cell transcriptomes, we identified 79 regulons with significant *cis*-regulatory motif enrichment showing significant changes in response to at least one activity-pattern (**Fig. 2d**, **Supplementary Fig. 5**). Many early-response ARGs encode TFs that regulate a subsequent wave of late-response gene expression. SCENIC analysis of newly-transcribed RNAs revealed neuronal activity-dependent increase in TF regulon activity of well-established IEG TFs (e.g., Fos, Jun, and Egr family of TFs) as well as constitutively expressed TFs such as Srf and Mef2, both of which are activated by multiple calcium-dependent signaling pathways and undergo post-translational modifications (e.g., phosphorylation) in response to neuronal activity (**Fig. 2d**). Thus, these TFs represent a group of TFs that are the main mediator of activity-dependent transcription. This group also included several TFs that have not been previously implicated in neuronal activation, including Maff and Hspa5, both of which exhibited higher expression and regulon activity upon activation (**Fig. 2d**). In addition, both Rfx3, a ciliogenic TF^19^, and the neural cell fate regulator Pou3f1^20^ were identified as potential novel regulators induced by neuronal activity. Furthermore, we observed regulon activity of cell-type-specific TFs (Neurod1/2 for Ex, Sox2/3 for Ex-NP and Dlx1/2 for Inh), which are associated with stable cell-type identities and detected in both newly-transcribed and pre-existing transcriptomes (**Fig. 2d**). Collectively, these data suggest that scNT-Seq allows for the analysis of cell-type-specific TF regulons that underly the dynamic cellular responses of both new and old transcriptomes to acute and sustained stimuli.

### scNT-Seq enables newly-synthesized transcripts based RNA velocity analysis

A fundamental question in gene regulation is how transcriptional states in single cells change over time in response to both acute (minutes) and sustained (hours to days) external stimuli. Recent work showed that the time derivative of the gene expression state, termed “RNA velocity,” can be estimated by distinguishing between unspliced (intronic reads) and spliced (exonic reads) mRNAs in scRNA-Seq datasets^21^. The RNA velocity measurement can predict the future state of individual cells on a timescale of hours. Because ultra-short metabolic labeling (5 minutes of 4sU pulse) can be used to identify transient RNAs^22^ and scNT-Seq enables direct measurements of new and old transcripts from the same cell, we next sought to augment the RNA velocity model to predict rapid cell state changes by replacing unspliced/spliced read levels with UMI-based newly-transcribed and total RNA levels. We focused on the excitatory neurons as this population of neurons robustly responds to neuronal activation. To compare unspliced/spliced ratio-based RNA velocity (termed “splicing RNA velocity”) and newly-transcribed RNA fraction-based velocity (termed “new RNA velocity”) directly, we projected and aligned both velocity fields onto the same UMAP plot (**Fig. 3a**). While splicing RNA velocity showed a steady directional flow (arrows) from 60 to 120 min (left in **Fig. 3a**), new RNA velocity revealed two distinct phases of velocity flows (early phase: 15 to 30 min; late phase: 60 to 120 min) (right in **Fig. 3a**). Thus, our analyses are consistent with recent reports that these two types of velocity measurements convey different but complementary information on the future state of a cell^8, 21^.

**Figure 3.**
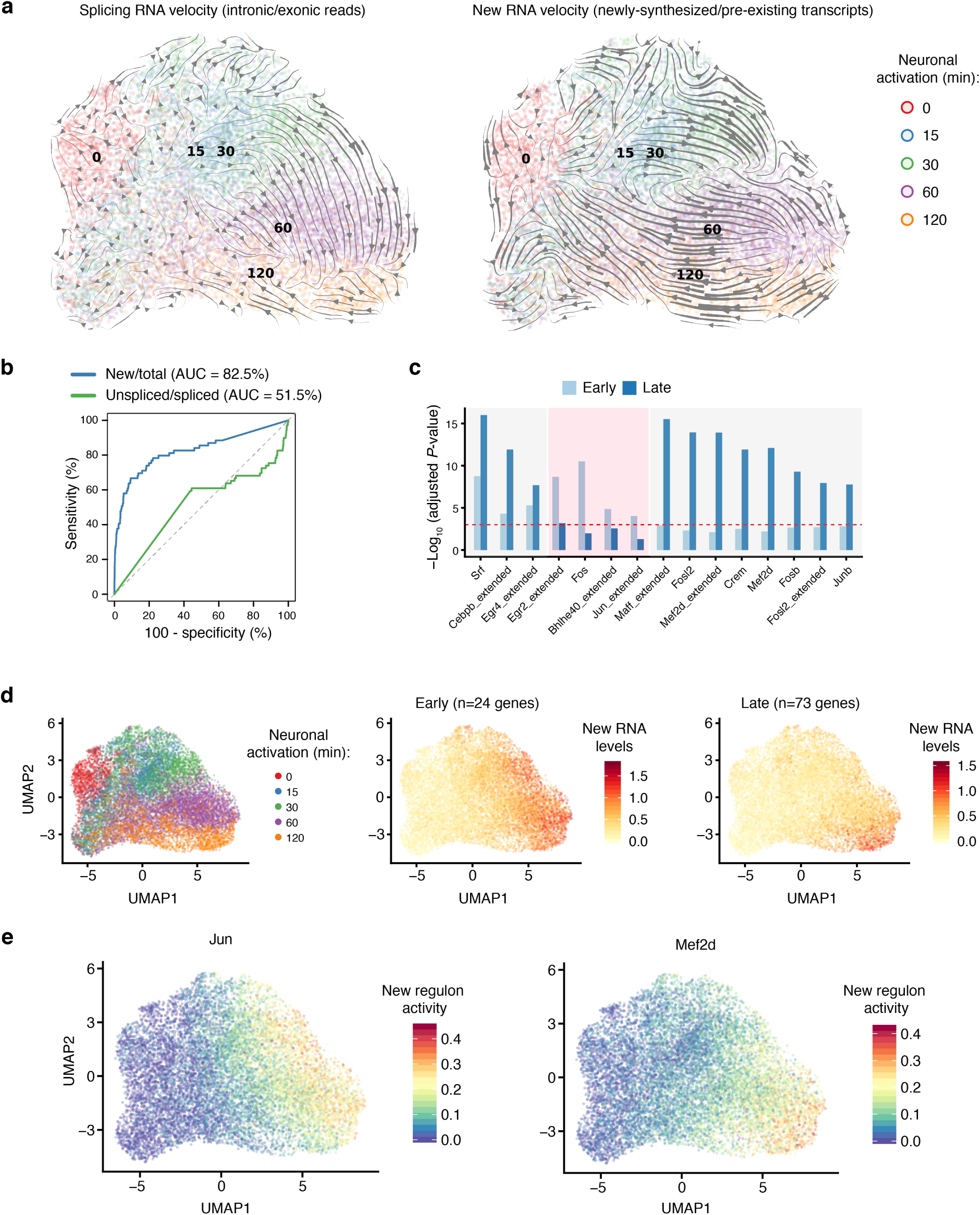
scNT-Seq reveals neuronal activity-induced new RNA velocity. (**a**) RNA velocity analysis of excitatory neurons using ratios of unspliced-to-spliced transcripts (splicing RNA velocity, left panel) or new-to-total transcripts (new RNA velocity, right panel) as inputs. The cells are color-codeed by different neuronal activity durations. The arrow indicates the projection from the observed state to extrapolated future state. (**b**) Comparison between splicing and new RNA velocity. Spliced counts or new RNA counts (30 min KCl stimulation) were used to predict induction of 137 known primary response genes^16^. The area under the curve (AUC) values of predictions from these two methods are shown. (**c**) Enrichment analysis of TF regulons for early- or late-response genes. Bar plots showing the significance of enriched TF regulons by hypergeometric test. TF regulons that are enriched for both groups, only early- or only late-response genes are highlighted by three boxes (left to right). The cut off of significance was indicated by red dash line (adjusted *P*-value < 0.001). (**d**) UMAP plots showing clustered Ex neurons from five time-points (left), expression level of early-response (middle), and late-response genes (right). (**e**) UMAP plots showing the regulon activity of two representative TFs regulating early- (Jun) or late-response genes (Mef2d), respectively. The UMAP plots in Fig. 3d-e are the same as in Fig. 3a.

New RNA velocity is grounded in direct measurements of newly-transcribed RNAs, and this approach promises to improve quantitative analysis of the dynamics of cell states, particularly for the analysis of transient and dynamic responses to rapid external stimulation. Indeed, new RNA velocity (area under the receiver operating characteristic curve (AUC) value = 82.5%) outperformed splicing RNA velocity (AUC=51.5%) analysis in predicting the activity-induced primary response genes (PRGs, 137 genes) that were identified in targeted bulk RNA-seq analysis of primary cortical neurons^16^ (**Fig. 3b**). These results show that new RNA velocity is more reliable in revealing the rapid transcriptional changes following acute neuronal activation.

To further explore the molecular basis underlying the two distinct phases of new RNA velocity flows, we identified the TF regulons that are significantly associated with early- (n=24; induced at 15 or 30 min) or late-response (n=73; induced at 60 or 120 min) genes (**Fig. 3c**, **Supplementary Fig. 4c**). We further projected aggregated newly-transcribed RNA levels of early- and late-response genes onto respective RNA velocity fields to visualize the relationship between distinct activity-patterns and ARG induction (**Fig. 3d**). Next, we projected TF regulon activity onto new RNA velocity field, revealing distinct activity patterns of TFs primarily associated with either early- (Jun) or late- (Mef2d) response genes in dynamic cell state changes (**Fig. 3e**). Together, these results indicate that integrative analysis of new RNA velocity and TF regulon activity can be a powerful approach to reveal molecular insights into dynamic cellular processes.

### scNT-Seq captures immediate transcription changes during the pluripotent-to-totipotent stem cell state transition

Determining temporal RNA dynamics such as immediate transcriptional events within rare, transient cell populations is critical to understanding cell state transition but has remained a significant challenge for stem cell research. Recent advances in multiplex single-molecule imaging using sequential fluorescence *in situ* hybridization (seqFISH) and intron probes allows for the detection of nascent transcription sites at the single-cell level^23^; however, such imaging approaches are currently time-consuming and have not been used to study rare stem cell populations at scale. Using mESC cultures as a model, we sought to determine whether scNT-Seq could be used to directly investigate immediate transcription changes and cell state transition within rare stem cell populations.

Cultured mESCs are derived from the inner cell mass of pre-implantation blastocysts and exhibit a high level of transcriptional heterogeneity^24^. Interestingly, cells resembling 2-cell-stage embryos (2C-like cells), which are in a totipotent state and can differentiate into both embryonic and extraembryonic tissues, arise spontaneously in mESC cultures^25^. Nearly all mESCs reversibly cycle between pluripotent and totipotent 2C-like state in culture. However, the 2C-like cells are rare in mESC cultures (<1% in standard culture conditions)^25^, making it challenging to study these totipotent cells with traditional methods. Recent studies using scRNA-Seq revealed changes in total RNAs during the transition to rare 2C-like state^26, 27^ and identified an intermediate state during this low-frequency transition process, supporting a stepwise model of transcriptional reprogramming of the pluripotent to 2C-like state transition^26, 27^. However, these studies cannot distinguish newly-transcribed from pre-existing RNAs at each state. Thus, a more detailed understanding of direct gene regulatory changes in response to state transition is lacking.

To capture newly-transcribed RNAs and characterize GRNs in different states, mESCs were metabolically labeled with 4sU for 4 hours and were subjected to scNT-Seq analysis (**Fig. 4a**). After quality filtering, we obtained 4,633 single-cell transcriptomes from two biological replicates (**Supplementary Fig. 6a, b**).

**Figure 4.**
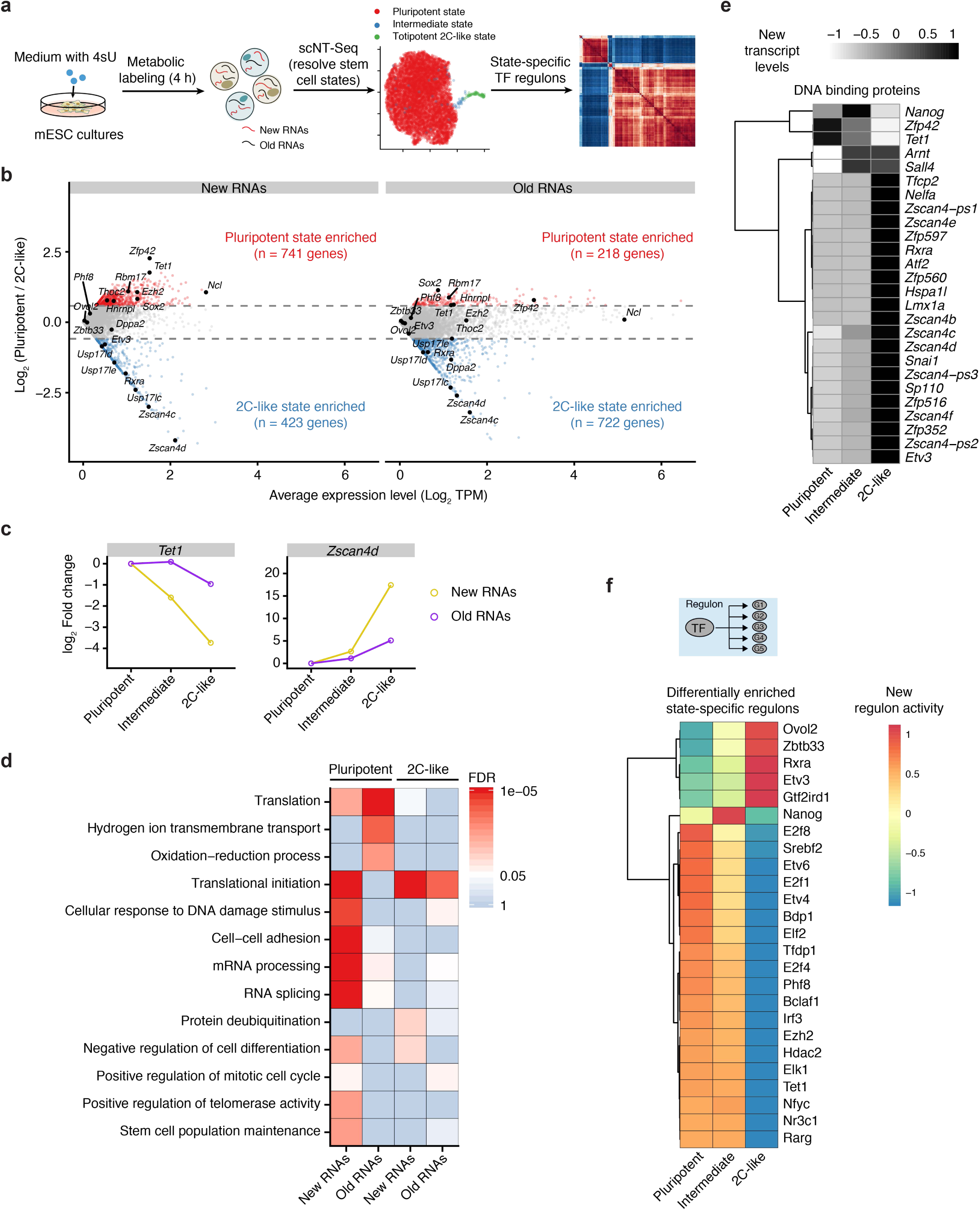
scNT-Seq captures both new and old transcripts in different states of mESCs. (**a**) Schematics of the scNT-Seq analysis of mESCs. (**b**) MA-plot showing differential gene expression of new and old transcripts between pluripotent and 2C-like states. Dashed line denotes 1.5-fold differences. (**c**) The line plots showing new and old transcript levels of *Tet1* and *Zscan4d* in pluripotent, intermediate, and 2C-like state. (**d**) Heat map showing enriched GO terms of state-specific genes. Significance of enrichment (FDR) is scaled by colors. (**e**) Heat map showing new transcript levels of state-specific genes encoding DNA-binding proteins. (**f**) Heat map showing the regulon activity of state-specific TFs or epigenetic regulators. The color scales depict the regulon activity.

Unbiased clustering not only readily separated contaminating mouse feeder cells (*Col1a2*/*Thbs1*+) from mESCs, but also identified three principal states within mESCs (**Supplementary Fig. 6a-c**). The majority of mESCs (98.3%) belong to the pluripotent state as they are positive for pluripotency-genes such as *Sox2* and *Zfp42* (also known as Rex1) but are negative for 2C-embryo-specific transcripts such as the *Zscan4* gene family (**Fig. 4b**). Approximately 0.7% of cells express high levels of 2C-like state markers, including *Zscan4* genes, *Zfp352* and *Usp17lc*, but low levels of *Sox2*, suggesting that this rare population contains predominantly totipotent 2C-like cells. These results are consistent with previous FACS analysis of 2C-reporter lines^25, 28^, indicating that scNT-Seq accurately captures this rare cell state. Interestingly, we also identified an intermediate state (∼1.0%) which expresses low levels of *Zscan4* genes, but high levels of *Nanog* and *Tbx3* (**Supplementary Fig. 6c, d**). Unlike previous studies which overexpressed a master regulator, Dux, to induce 2C-like state^27^, we did not observe a significant decrease in *Sox2* expression in the intermediate state (**Supplementary Fig. 6c**), highlighting the potential difference between the Dux-induced 2C-transitions and the naturally occurring pluripotent-to-totipotent transition process.

Comparative analysis of new and old transcriptomes revealed many state-specific genes (e.g., *Sox2* and *Zscan4d*) are associated with a higher proportion of new transcripts (consistent with their regulatory roles), whereas house-keeping genes, such as *Gapdh*, are of slower turnover rate (**Supplemental Fig. 7a, b**). For many state-specific genes (e.g., *Zfp42* and *Zscan4d*), their new RNA levels exhibited a more pronounced difference between states than the change of pre-existing RNAs (**Fig. 4b, c**). Gene Ontology (GO) enrichment analysis further revealed that state-specific new and old transcripts were enriched for different GO terms (**Fig. 4d**). For instance, many pluripotent-state-specific new RNAs (but not old RNAs) are functionally related to post-transcriptional regulatory processes, including translational initiation, mRNA processing, and RNA splicing. Furthermore, 2C-state-specific new transcripts are preferentially enriched for deubiquitinating enzyme-related genes such as *Usp17lc/d/e* (**Fig. 4b, d**). These results underscore that newly-synthesized RNAs are more robust than steady-state transcripts to uncover genes related to state-specific biological processes within heterogeneous stem cell populations.

DNA-binding proteins, such as TFs and epigenetic regulators, show rapid gene expression changes during pluripotent-to-2C transition^27, 29^. Our analysis of immediate transcription changes during the pluripotent-to-2C transition identified 26 genes encoding DNA-binding proteins that showed a significant difference at new RNA levels between states (**Fig. 4e**). Consistent with previous scRNA-Seq analysis of sorted cell populations^29^, 13 out of 14 commonly detected genes (*Tet1*, *Sox2*, *Nanog*, *Rex1*/*Zfp42*, *Sp110*, *Zfp352,* and *Zscan4* family) showed similar patterns in our scNT-Seq analysis of new RNAs (**Fig. 4e**). To further investigate TF/epigenetic regulator activity during the pluripotent-to-2C transition, we applied SCENIC analysis to both new and old transcriptomes. Among all regulons uncovered in mESCs (**Supplementary Fig. 8a**), we identified 25 TF/epigenetic regulators showing differential new regulon activity between states (**Fig. 4f** and **Supplementary Fig. 8b**). The activity of several positive regulators of the cell cycle, such as the E2F family of TFs, decreased in 2C-like states, which is consistent with previous observations that 2C-like cells are associated with longer cell cycles^30^. Interesting, new RNA levels and regulon activity of Nanog peak in intermediate state compared to pluripotent and totipotent 2C-like states, suggesting that this TF may play an unrecognized regulatory role in intermediate state. In addition, several epigenetic regulators, including Phf8, Hdac2, Ezh2, and Tet1, were associated with a decrease in regulon activity (**Fig. 4f**), suggesting a role of these enzymes in promoting the pluripotent state. Notably, we did not identify the regulon activity of Zscan4 in 2C-like cells, potentially due to the lack of Zscan4 motif information in the SCENIC TF database. Together, scNT-Seq directly captures state-specific newly-transcribed RNAs in rare totipotent cells in heterogeneous mESC cultures and allows analysis of dynamics of new regulon activity during state transitions.

### TET-dependent regulation of the stepwise pluripotent-to-2C transition

The TET family of proteins (Tet1-3) are DNA dioxygenases that mediate active DNA demethylation at many *cis*-regulatory elements such as promoters and distal enhancers^31, 32^, thereby regulating gene expression. During pluripotent-to-2C transition, both new RNA level and regulon activity of *Tet1* rapidly decreased (**Fig. 4c, f**). The new transcript level of *Tet2* also decreased in both intermediate and 2C-like states, while *Tet3* was undetected in all three states (**Supplementary Fig. 7c**). Interestingly, genetic inactivation of all TET proteins in mESCs resulted in a substantially higher proportion of 2C-like cells, and elevated expression level of 2C-specific transcripts^28^.

To better understand the role of TET enzymes in state transition, we generated mESCs deficient for all three Tet proteins (*Tet1/2/3* triple knockout, *Tet*-TKO) via CRISPR/Cas9 genome editing using previously-tested sgRNAs^33^ and analyzed isogenic WT and *Tet*-TKO mESCs (J1 strain) by scNT-Seq after 4 hours of metabolic labeling. The genotype of *Tet*-TKO cells was confirmed by both Sanger sequencing and reads from scNT-Seq (**Supplementary Fig. 9a**). Clustering and new RNA velocity analyses of combined WT and *Tet*-TKO mESCs revealed two discrete phases of new RNA velocity flows (arrows). The velocity flow within the pluripotent state is potentially driven by cell-cycle progression (**Supplementary Fig. 9b, c**), whereas new RNA velocity field of intermediate/2C-like states exhibited a strong directional flow from intermediate to 2C-like state (**Fig. 5a, b**). These observations are consistent with a stepwise model of the state-transition process and suggest that the pluripotent-to-intermediate state transition is a rate-limiting step.

**Figure 5.**
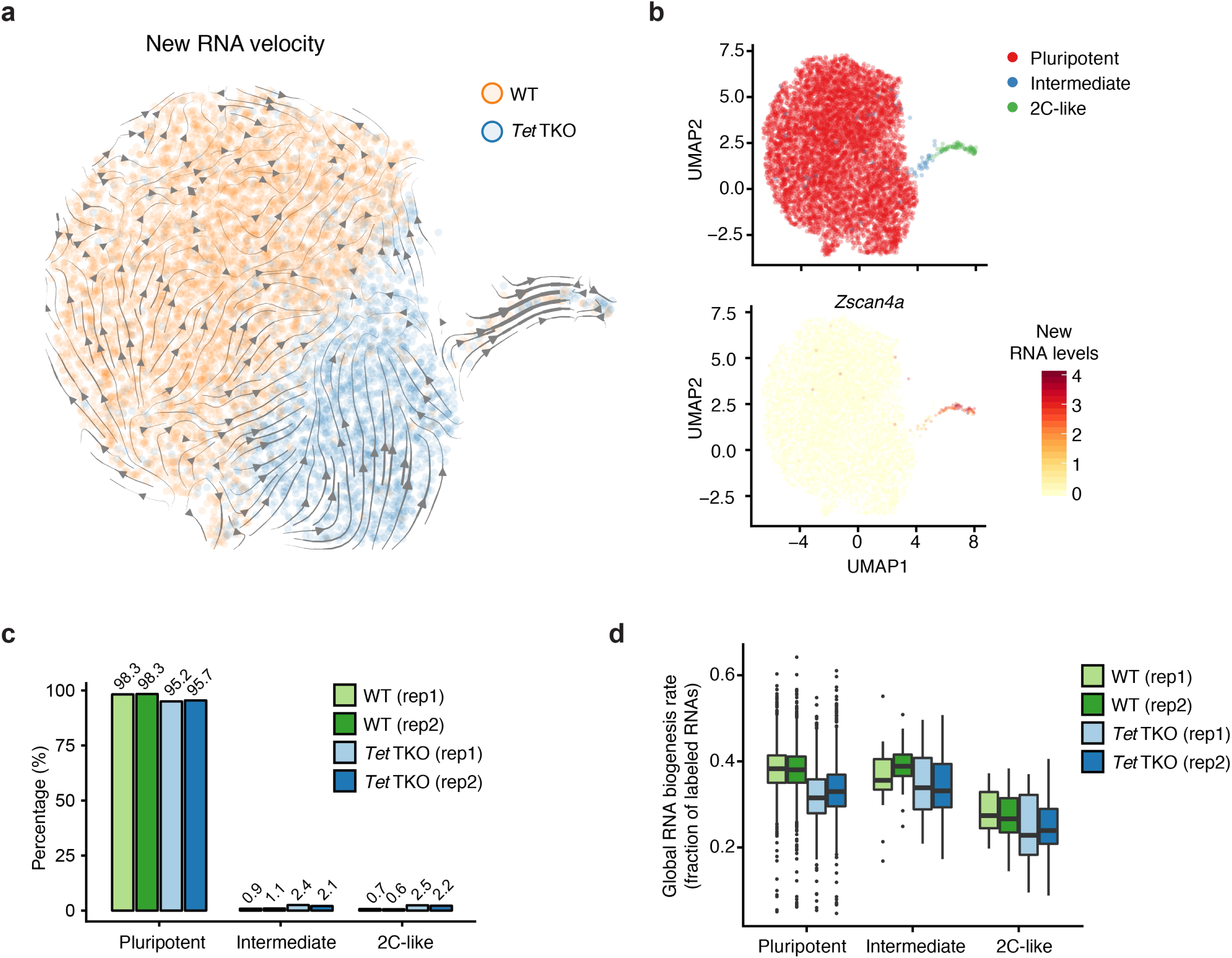
scNT-Seq reveals TET-dependent regulation of the plutipotent-to-2C transition. (*a*) New RNA velocity analysis of combined wild-type (WT) and *Tet*-TKO mESCs. The arrow indicates the projection from the observed state to extrapolated future state. (**b**) Projection of cell state and the new RNA level of the *Zscan4a* gene on the same UMAP plot as in Fig. 5a. (**c**) Relative composition of three stem cell states in WT (4,633 cells) and *Tet*-TKO (2,319 cells) mESCs. (**d**) The fraction of labeled transcripts in WT and *Tet*-TKO mESCs across three states. The box plots display the median (center line) and interquartile range (IQR, from the 25th to 75th percentile), the whiskers represent 1.5 times the interquartile range, and the circles represent outliers.

WT and *Tet*-TKO cells were separately clustered within the pluripotent state and *Tet*-TKO (but not WT) cells were found immediately next to the intermediate/2C-like states (**Fig. 5a, b**), suggesting that *Tet*-TKO cells are transcriptionally reprogrammed to adopt a more poised state to transition to intermediate/2C-like states. Compared to WT cells, *Tet*-TKO cells exhibited a 2.2-fold or 3.6-fold increase in intermediate or 2C-like states (**Fig. 5c**), respectively, indicating that Tet enzymes negatively regulate the transition from pluripotent to intermediate/2C-like state in WT cells. Because pluripotent and 2C-like cells are globally distinct in levels of both major histone modifications^25^ and DNA methylation^26^, we next examined global levels of newly-synthesized transcripts in each state. Interestingly, aggregating new transcriptomes within specific states revealed that a totipotent 2C-like state is associated with markedly lower levels of global RNA biogenesis compared to pluripotent and intermediate states in WT cells (**Fig. 5d**). In contrast, *Tet*-TKO cells already exhibited a substantially lower level of global transcription than WT cells in pluripotent state (**Fig. 5d**), suggesting that Tet proteins may act as an epigenetic barrier for the pluripotent-to-2C transition by maintaining a pluripotent state-specific RNA biogenesis profile.

### Pulse-chase scNT-Seq enables transcriptome-wide measurement of mRNA stability in rare 2C-like cells

Pulse-chase assays combined with bulk SLAM-Seq have been used to measure mRNA stability in mESCs^5^, but rare intermediate and 2C-like cells have not been studied. Given the enhanced sensitivity of scNT-Seq compared to bulk assays, combining a pulse-chase strategy with scNT-Seq may enable transcriptome-wide measurement of mRNA stability in rare cell populations. To test this, we metabolically labeled mESC cultures with 4sU for 24 hours (pulse), followed by a washout and chase using medium containing a higher concentration of uridine. Cells from multiple chase time-points were harvested and cryo-preserved first, and all samples were then re-hydrated and simultaneously analyzed by scNT-Seq analysis. After computing the proportion of labeled transcripts for each gene at every time-point relative to 0 h (right after pulse), we calculated the half-life (t1/2) of mRNAs by fitting a single-exponential decay model in each cell state (**Fig. 6a**). In total, 20,190 cells were profiled by scNT-Seq from 7 time-points (**Fig. 6b**). After 24 hours of metabolic labeling, we observed a substantial accumulation of T-to-C substitution (**Fig. 6c**), which is consistent with bulk assay observations^5^. The T-to-C substitution rate decreased over time and returned to the baseline level 24 hours after chase (**Fig. 6c**). Inspection of two genes, *Sox2* and *Topa2a*, revealed that total RNA levels did not change across the 24-hour time course, whereas the level of metabolically labeled transcripts decreased over time (**Supplementary Fig. 10a**), confirming that this strategy can accurately measure mRNA decay in a transcript-specific manner.

**Figure 6.**
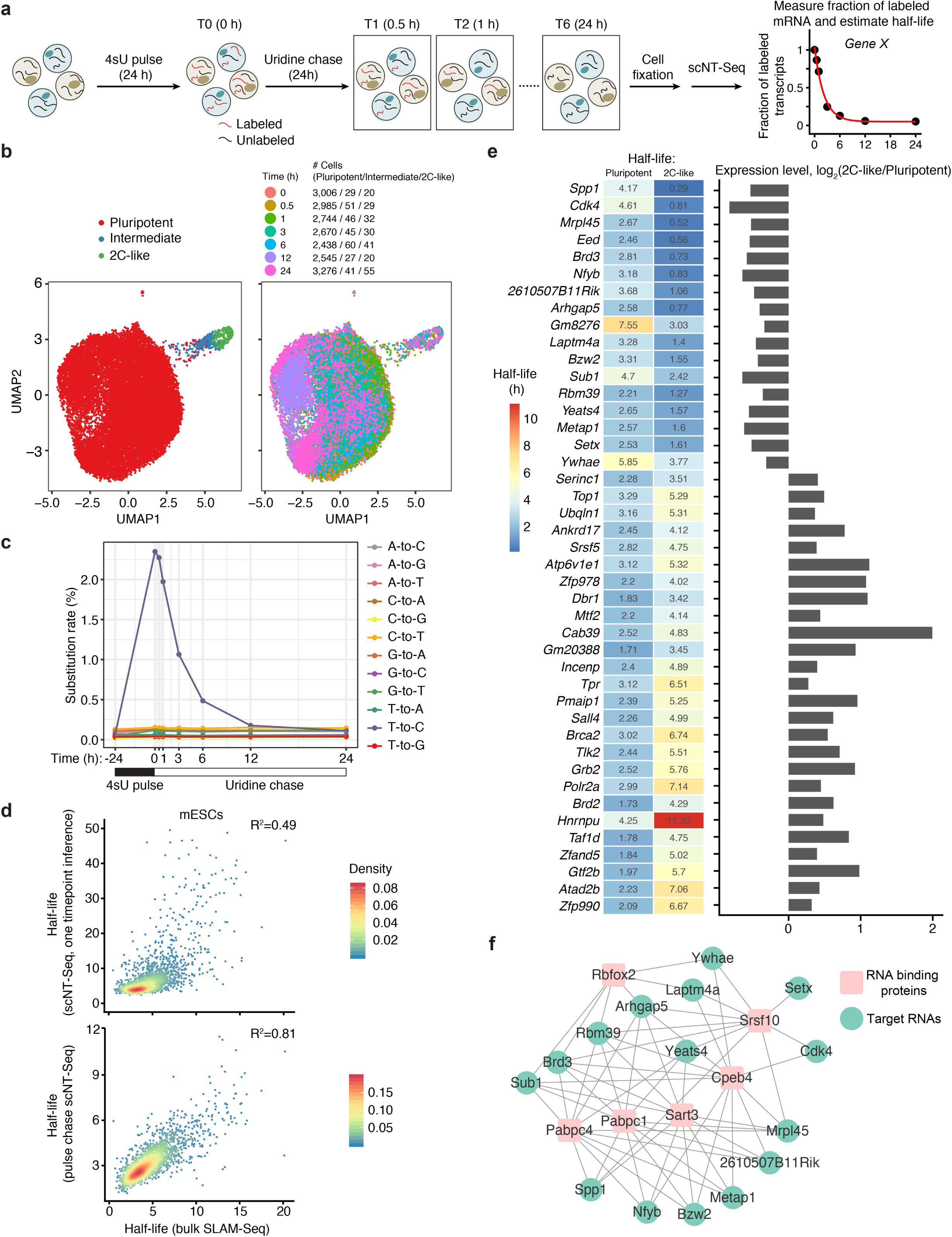
Pulse-chase scNT-Seq reveals cell state-specific mRNA decay in mESCs. (**a**) Schematics of pulse-chase scNT-Seq experiment. (**b**) UMAP plots of 20,190 mESCs profiled in the pulse-chase scNT-Seq analysis. Cells are colored by three states (left) or by 7 time-points (right). Cell numbers of each state across different time-points are also shown. (**c**) Line plots showing temporal changes of nucleotide substitution rates. (**d**) Scatterplots showing the correlation of RNA half-life measurements between this study (top: one timepoint inference analysis; bottom: pulse-chase analysis) and bulk SLAM-Seq^5^. (**e**) Heat map (left) and bar plot (right) showing concordant change of RNA half-life (left) and gene expression (right) between pluripotent and 2C-like states, respectively. (**f**) Network of RNA-binding proteins (pink quadrate) and their binding targets (green circle). Enriched RNA-binding proteins were inferred by motif sequenced enriched in 3’UTR of target genes.

Based on expression of marker genes, clustering analysis readily separated mESCs into three states (n=7, pluripotent: 97.4%+/-0.78%, intermediate: 1.5%+/-0.48%, and 2C-like: 1.1%+/-0.38%) (**Fig. 6b** and **Supplementary Fig 10b**), suggesting that this pulse-chase approach did not significantly alter the state transition. By filtering out genes expressed in less than 5% of cells, we were able to determine the half-life of 2,310 genes in pluripotent state, and the RNA half-life determined by pulse-chase scNT-Seq is highly concordant with previous observation derived from bulk SLAM-Seq assays^5^ (R^2^=0.81, **Fig. 6d**). RNA stability can also be estimated by assuming simple exponential kinetics, which is calculated by measuring the ratio of labeled and unlabeled transcripts after metabolic labeling for a specific time^34^. However, RNA half-life estimated from one timepoint labeling (4sU, 4 hour) experiment is substantially less correlated with measurements derived from bulk assays (R^2^=0.49, **Fig. 6d**). These results suggest that a pulse-chase strategy may more reliably measure RNA decay rate. Furthermore, the top 10% most stable and unstable transcripts were enriched for similar GO terms that are uncovered by bulk SLAM-Seq assays^5^ (**Supplementary Fig. 10c**). Further analysis of cells in intermediate and 2C-like states revealed the half-life of 1,743 and 821 transcripts in these rare cell states, respectively. Next, we analyzed commonly detected transcripts between cell states to reveal state-specific regulation of mRNA stability (**Supplementary Fig. 11**). Thus, scNT-seq enables transcriptome-wide measurement of RNA stability in rare stem cell populations in heterogeneous mESC cultures.

Because dynamic changes in RNA levels are regulated by the interplay of RNA biogenesis and degradation, we sought to investigate the relationship between mRNA stability and gene expression during the pluripotent-to-2C transition (**Supplementary Fig. 12**). Consistent with previous findings^2^, RNA stability and total mRNA levels are not highly correlated, suggesting mRNA stability is not the major contributor to total RNA level during stem cell state transition. Nevertheless, we identified a group of genes showing coordinated changes of RNA stability and gene expression level between pluripotent and 2C-like states, suggesting mRNA stability may play a role in regulating total mRNA level for a subset of genes (**Fig. 6e**). Further analysis showed that six RNA-binding protein (RBP) binding motifs were enriched in 3’UTR of 15 genes showing coordinated changes (**Fig. 6f**). Among these RBPs, the Pabpc family proteins (Pabpc1 and Pabpc4) are known regulators of mRNA stability and translation efficiency by binding poly-A tails of mRNAs^35^. Collectively, these results indicate that scNT-Seq can serve as a powerful approach to investigate RNA stability and post-transcriptional regulatory mechanisms in rare cells within a heterogeneous population of cells.

## DISCUSSION

By combining the widely accessible Drop-Seq platform and TimeLapse chemistry, scNT-Seq provides a novel strategy for high-throughput analysis of new and old transcriptomes from the same cell. A major advantage of scNT-Seq is its ability to accurately estimate newly-transcribed RNA levels using UMI-based statistical correction, which not only reveals acute transcriptional changes that are not apparent in standard scRNA-seq, but also allows for the analysis of dynamic TF regulatory networks and single-cell transcriptional trajectory. Our scNT-Seq is conceptually similar to sci-fate^36^, a recently developed method that integrates SLAM-Seq chemistry with single-cell combinatorial indexing RNA-Seq. While scNT-Seq directly captures whole-cell total RNAs on barcoded beads in nanoliter droplets, sci-fate requires paraformaldehyde fixation and permeabilization of cells for *in situ* chemical conversion and reverse transcription, which may pose a challenge to unbiasedly recover both nuclear (enriched for newly-transcribed) and cytoplasmic (enriched for pre-existing) RNAs for accurately estimating new RNA fractions.

Because the purine analog, 6-thioguanine (6tG), enables G-to-A conversions by TimeLapse chemistry in bulk RNA-Seq^37^, dual labeling of 4sU and 6tG followed by scNT-Seq may allow two independent recordings of newly-synthesized transcriptome at single-cell levels. With new computational approaches such as Dynamo^38^, which can take advantage of controlled metabolic labeling to predict past and future cell states over an extended time period, high-throughput single-cell transcriptomic analysis of newly-transcribed and pre-existing RNAs can open new lines of inquiry regarding cell-type-specific RNA regulatory mechanisms and provide a broadly applicable strategy to investigate dynamic biological systems.

## ACKNOWLEDGEMENTS

We are grateful to all members of the Wu lab for helpful discussion. This work was supported by the Penn Epigenetics Institute, the National Human Genome Research Institute (NHGRI) grants R00-HG007982 and R01-HG010646, National Heart Lung and Blood Institute (NHLBI) grant DP2-HL142044, and National Cancer Institute (NCI) grant U2C-CA233285 (to H.W.). The work of P.C. is partially supported by Stand-Up-To-Cancer Convergence 2.0.

## AUTHOR CONTRIBUTIONS

H.W., Q.Q., and P.H. conceived of and developed the scNT-Seq approach. Q.Q. conducted most of the experiments. P.H. generated *Tet*1/2/3 TKO mESC line and performed most of the computational analysis. K.G. and P.C. contributed to the statistical modeling of T-to-C substitution. Q.Q., P.H., and H.W. analyzed the results and wrote the manuscript, with contributions from all of the authors. H.W. supervised the project.

## Online Methods

### Experimental procedures

#### Mouse embryonic stem cell cultures and 4sU labeling

*Tet*1/2/3 triple-knock out (*Tet*-TKO) J1 mESCs were generated by CRISPR/Cas9 genome editing as previously described^33^ and the genotype was confirmed by Sanger sequencing and scRNA-Seq. Wild-type (WT) and *Tet*-TKO J1 mESCs (ATCC, SCRC-1010) were initially cultured in presence of Mitomycin C inactivated mouse embryonic fibroblasts on 0.1% gelatin-coated (Millipore, ES-006-B) 6-well plates in Dulbecco’s Modified Eagle’s Medium (DMEM) (Gibco, 11965084) supplemented with 15% fetal bovine serum (Gibco, 16000044), 0.1 mM nonessential amino acid (Gibco, 11140050), 1 mM sodium pyruvate (Gibco, 11360070), 2 mM L-glutamine (Gibco, 25030081), 50 μM 2-mercaptoethanol (Gibco, 31350010), 1 μM MEK inhibitor PD0325901 (Axon Med Chem, Axon 2128) and 3 μM GSK3 inhibitor CHIR99021 (Axon Med Chem, Axon 2128), and 1,000 U/mL LIF (Gemini Bio-Products, 400-495-7).

4-thiouridine (4sU) (Alfa Aesar, J60679) were dissolved in DMSO to make 1 M stock. Before 4sU labeling experiments, WT and *Tet*-TKO mESCs were passaged and cultured in feeder-free conditions (0.1% gelatin-coated plates) for 48 hrs. For 4sU labeling in WT and Tet TKO mESC cultures, the medium was replaced with fresh mESC medium supplemented with 100 μM 4sU. After 4 h, mESCs were rinsed once with PBS and dissociated with TrypLE-Express (Gibco, 12605010) for 5 min at 37°C. After neutralizing with culture medium, cells were pelleted at 1,000 rpm for 3 min. After cell counting with the Countess II system, the single cell suspension was diluted to 100 cells/uL with DPBS containing 0.01% BSA for scNT-Seq analysis.

#### Human K562 cell cultures and species mixing experiments

Human K562 cells (ATCC, CCL-243) were cultured in RPMI media supplemented with 10% FBS (Sigma, F6178). For species mixing experiments, the mESC or K562 media was replaced with media supplemented with 4sU (100 μM). After 4 h, the mESCs and K562 cells were rinsed with PBS and harvested for scNT-Seq analysis.

#### Mouse primary neuronal culture and stimulation

Mouse cortices were dissected from embryonic day 16 (E16) C57BL/6 embryos of mixed sex (Charles River). Cortical neurons were dissociated with papain (Worthington) and plated on 6-well plates (at a density of ∼600,000 cells/well) coated with poly-ornithine (30mg/mL, Sigma, P2533). Mouse cortical neuronal cultures were maintained in neurobasal media (Gibco, 21103049) supplemented with B27 (Gibco, 17504044), 2 mM GlutaMAX (Gibco, 35050061), and 1X Penicillin/streptomycin (Gibco, 15140122).

After 4 days *in vitro* culture, primary cortical cultures were stimulated with a final concentration of 55mM potassium chloride (KCl) for various time (0, 15, 30, 60, and 120 minutes). Before neuronal activation, the fresh media supplemented with 4sU (200 μM) was added. After 4sU labeling (2 h), the cells were washed with PBS once and were digested with 0.05% Trypsin-EDTA (Gibco, 25300054) for 20 min at 37°C, then replaced the buffer with 1 mL of DPBS and dissociated cells with a cell-scraper. After cell counting with Countess II system, the single cell suspension was diluted to 100 cells/μL with DPBS containing 0.01% BSA for scNT-Seq analysis.

#### Cell fixation, cryopreservation and rehydration for scNT-Seq

The cell fixation was performed as previously described^39^. Cultured mESCs in 6-well plates were digested with TrypLE-Express and harvested as aforementioned. After washing once with DPBS, the cells were resuspended with 0.4 mL of DPBS containing 0.01% BSA. Split the cells to two 1.5 mL LoBind tubes (Eppendorf) and add 0.8 mL methanol dropwise at final concentration of 80% methanol in DPBS. After mixing and incubating the cell suspension on ice for 1 hour, store the fixed cells in LoBind tubes at -80°C freezer for up to one month. For rehydration, cells were kept on ice after moved from –80 °C and kept in the cold throughout the procedure. After the cells were spun-down at 1,000 g for 5 min at 4°C, Methanol-PBS solution was removed and cells were resuspended in 1 mL 0.01% BSA in DPBS supplemented with 0.5% RNase-inhibitor (Lucigen, 30281-2). After cell counting with the Countess II system, the single cell suspension was diluted to 100 cells/μL and immediately used for scNT-Seq analysis.

#### Pulse-chase experiment for RNA stability analysis

Remove the medium from plates and add mESC medium supplemented with 200 μM 4sU^5^. The fresh medium with 4sU were changed every 4 h to enhance 4sU incorporation and mESCs were labeled for 24 hrs. After 4sU-labeling, the 4sU-containing medium was removed and cells were washed once with DPBS. Then mESC medium containing 10 mM Uridine (Sigma, U6381) was added to the culture before cells were harvested at different time-points ranging from 0.5 hour to 24 hours. Cells were fixed with methanol as aforementioned and store at -80°C for future use.

#### scNT-Seq library preparation and sequencing

Drop-Seq was performed as previously described with minor modifications^10^. Specifically, the single cell suspension was diluted to a concentration of 100 cells/μL in DPBS containing 0.01% BSA. Approximately 1.5 mL of diluted single cell suspension was loaded for each scNT-Seq run. The single-nucleus suspension was then co-encapsulated with barcoded beads (ChemGenes) using an Aquapel-coated PDMS microfluidic device (uFluidix) connected to syringe pumps (KD Scientific) via polyethylene tubing with an inner diameter of 0.38 mm (Scientific Commodities). Barcoded beads were resuspended in lysis buffer (200 mM Tris-HCl pH8.0, 20 mM EDTA, 6% Ficoll PM-400 (GE Healthcare/Fisher Scientific), 0.2% Sarkosyl (Sigma-Aldrich), and 50 mM DTT (Fermentas; freshly made on the day of run) at a concentration of 120 beads/μL. The flow rates for cells and beads were set to 3,200 μL/hour, while QX200 droplet generation oil (Bio-rad) was run at 12,500 μL/h. A typical run lasts ∼20 min.

Droplet breakage with Perfluoro-1-octanol (Sigma-Aldrich). After droplet breakage, the beads were treated with TimeLapse chemistry to convert 4sU to cytidine-analog^6^. Briefly, 50,000-100,000 beads were washed once with 450 μL washing buffer (1 mM EDTA, 100 mM sodium acetate (pH 5.2)), then the beads were resuspended with a mixture of TFEA (600 mM), EDTA (1 mM) and sodium acetate (pH 5.2, 100 mM) in water. A solution of 192 mM NaIO4 was then added (final concentration: 10 mM) and incubated at 45°C for 1 hour with rotation. The beads were washed once with 1 mL TE, then incubated in 0.5 mL 1 X Reducing Buffer (10 mM DTT, 100 mM NaCl, 10 mM Tris pH 7.4, 1 mM EDTA) at 37°C for 30 min with rotation, followed by washing once with 0.3 mL 2X RT-buffer.

Reverse transcription and exonuclease I treatment were performed, as previously described, with minor modifications^10^. Specifically, up to 120,000 beads, 200 μL of reverse transcription (RT) mix (1x Maxima RT buffer (ThermoFisher), 4% Ficoll PM-400, 1 mM dNTPs (Clontech), 1 U/μL RNase inhibitor, 2.5 μM Template Switch Oligo (TSO: AAGCAGTGGTATCAACGCAGAGTGAATrGrGrG)^10^, and 10 U/ μL Maxima H Minus Reverse Transcriptase (ThermoFisher)) were added. The RT reaction was incubated at room temperature for 30 minutes, followed by incubation at 42°C for 120 minutes. To determine an optimal number of PCR cycles for amplification of cDNA, an aliquot of 6,000 beads (corresponding to ∼100 nuclei) was amplified by PCR in a volume of 50 μL (25 μL of 2x KAPA HiFi hotstart readymix (KAPA biosystems), 0.4 μL of 100 μM TSO-PCR primer (AAGCAGTGGTATCAACGCAGAGT^10^, 24.6 μL of nuclease-free water) with the following thermal cycling parameter (95°C for 3 min; 4 cycles of 98°C for 20 sec, 65°C for 45 sec, 72°C for 3 min; 9 cycles of 98°C for 20 sec, 67°C for 45 sec, 72°C for 3 min; 72°C for 5 min, hold at 4°C). After two rounds of purification with 0.6x SPRISelect beads (Beckman Coulter), amplified cDNA was eluted with 10 μL of water. 10% of amplified cDNA was used to perform real-time PCR analysis (1 μL of purified cDNA, 0.2 μL of 25 μM TSO-PCR primer, 5 μL of 2x KAPA FAST qPCR readymix, and 3.8 μL of water) to determine the additional number of PCR cycles needed for optimal cDNA amplification (Applied Biosystems QuantStudio 7 Flex). We then prepared PCR reactions per total number of barcoded beads collected for each scNT-Seq run, adding 6,000 beads per PCR tube, and ran the aforementioned program to enrich the cDNA for 4 + 10 to 12 cycles. We then tagmented cDNA using the Nextera XT DNA sample preparation kit (Illumina, cat# FC-131-1096), starting with 550 pg of cDNA pooled in equal amounts, from all PCR reactions for a given run. Following cDNA tagmentation, we further amplified the library with 12 enrichment cycles using the Illumina Nextera XT i7 primers along with the P5-TSO hybrid primer (AATGATACGGCGACCACCGAGATCTACACGCCTGTCCGCGGA AGCAGTGGTATCAACGCAGAGT*A*C)^10^. After quality control analysis using a Bioanalyzer (Agilent), libraries were sequenced on an Illumina NextSeq 500 instrument using the 75- or 150-cycle High Output v2 or v2.5 Kit (Illumina). We loaded the library at 2.0 pM and added Custom Read1 Primer (GCCTGTCCGCGGAAGCAGTGGTATCAACGCAGAGTAC) at 0.3 μM to position 7 of the reagent cartridge. Paired-end sequencing were performed on Illumina NextSeq 500 sequencer as described previously^11^. The sequencing configuration was 20 bp (Read1), 8 bp (Index1), and 60 or 130 bp (Read2).

#### SLAM-Seq reaction on barcoded Drop-Seq beads

After droplet breakage, the beads were washed once with NaPO4 buffer with 30% DMSO (50 mM, pH 8.0), then incubate beads in 500 μL reaction-mix containing 10 mM IAA for either 15 min at 50°C (standard SLAM-Seq) or 1 hour at 45°C (modified condition)^5^. Stop reaction by adding 10 μL 1 M DTT (final concentration: 20 mM).

### Bioinformatic analysis

#### Read mapping and quantification of labeled and unlabeled transcripts

Paired-end sequencing reads of scNT-Seq were processed as previous described^11^ with some modifications. Briefly, each mRNA read (read2) was tagged with the cell barcode (bases 1 to 12 of read 1) and unique molecular identifier (UMI, bases 13 to 20 of read 1), trimmed of sequencing adaptors and poly-A sequences, and aligned using STAR v 2.5.2a to the mouse (mm10, Gencode release vM13), human genome (GRCh38, Gencode release v23), or a concatenation of the mouse and human (for the species mixing experiment) reference genome assembly. Both exonic and intronic reads mapped to predicted strands of annotated genes were retained for the downstream analysis. To qualify the labeled and unlabeled transcripts, uniquely mapped reads with mapping score > 10 were grouped by UMI barcodes in every cell and were used to determine the T > C substitution using sam2tsv^40^, T > C substitutions with a base quality of Phred score > 27 were retained. For each experiment, locus with background T > C substitutions (detected in the sample without TFEA/NaIO4 treatment) was determined and was excluded for T > C substitution identification. After background correction of T > C substitution, a UMI was defined as labeled if there is a T > C substitution in any one of the reads belongs to that particular UMI. To this end, every UMI will be assigned to labeled or unlabeled based on the existence of T > C substitution (**Figs 1d and 5d**). For each transcript, the total number of uniquely labeled and unlabeled UMI sequences were counted and finally were assembled into a matrix using gene name as rows and cell barcode as columns.

#### Cell type clustering

For mouse cortical neurons and RNA-decay experiment (**Figs 2b** and **6b**), the raw digital expression matrices of new and old UMI counts were added up and loaded into the R package Seurat^41^ (v 2.3.4). For normalization, UMI counts for all cells were scaled by library size (total UMI counts), multiplied by 10,000 and transformed to log space. Only genes detected in >10 cells were retained. Cell with a relatively high percentage of UMIs mapped to mitochondrial genes were discarded (QC metrics in **Supplementary Table 1**). Moreover, cells with lower or higher detected genes were discarded (QC metrics in **Supplementary Table 1**). As a result, 20,547 cells of mouse cortical culture and 20,190 cells of mESC (in pulse-chase assay) were retained, respectively. The highly variable genes (HVGs) were identified using the function *FindVariableGenes* with the parameters: *x.low.cutoff* = .05,*y.cutoff* = .5 in Seurat, resulting in 2,290 HVGs of primary cortical culture sample and 2,165 HVGs of mESC sample. The expression level of highly variable genes in the cells was scaled and centered along each gene and was conducted to principal component analysis (PCA). The most significant 30 PCs were selected and used for 2-dimension reduction by uniform manifold approximation and projection^42^ (UMAP), implemented by the Seurat software with the default parameters. Clusters were identified using the function *FindCluster* in Seurat with the resolution parameter set to 1. To obtain high level of cell type classification, we merge the adjacent clusters in UMAP which highly expressed excitatory markers (*Neurod2* and *Neurod6*) and define it as “Ex” cluster in mouse cortical neurons, while several close clusters highly expressed *Sox2* were combined to “pluripotent” cluster in mESC Pulse-chase experiment. Cell type specific markers were identified using function *FindMarkers* in Seurat with wilcoxon rank sum test with default parameters.

Cell clustering will be affected both by genotype^28^ and by cell type. To enable directly comparative analyses within cell types between WT and *Tet*1/2/3 triple-knock out (*Tet*-TKO) mESCs, we used Seurat 3 (v. 3.0.0.9000) which was demonstrated as an effective strategy to perform joint analyses^43^ (**Supplementary Fig. 6a**). The raw digital expression matrices of new and old UMI counts were added up and loaded into the Seurat 3. For normalization, UMI counts for all cells were scaled by library size (total UMI counts), multiplied by 10,000 and transformed to log space. Only genes detected in >10 cells were retained. Cell with a relatively high percentage of UMIs mapped to mitochondrial genes (>=0.05) were discarded. Moreover, cells with fewer than 500 or more than 5,000 detected genes were discarded, resulting in 4,633 WT cells and 2,319 *Tet*-TKO cells. Top 2,000 HVGs were identified using the function *FindVariableFeatures* with “vst” method. Canonical correlation analysis (CCA) was used to identify common sources of variation between WT and *Tet*-TKO cells. The first 20 dimensions of the CCA was chosen to integrate the two datasets. After integration, the expression level of HVGs in the cells was scaled and centered along each gene and was conducted to PCA analysis. The 20 most significant PCs were selected and used for 2-dimension reduction by UMAP. Clusters were identified using the function *FindCluster* in Seurat with the resolution parameter set to 3. As above mentioned, adjacent clusters highly expressed *Sox2* were combined to “pluripotent” cluster.

#### Estimation of the fraction of newly-synthesized transcripts

We implemented a statistical modeling strategy as previously described with some modifications for UMI-based scNT-Seq analysis^6^. For each experiment, a binomial mixture model, was used to approximate the number of T-to-C substitutions *y*_*i*_ for each gene transcript *i*:

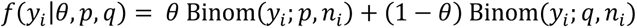

In this expression, *θ* is the fraction of new transcripts in each experiment, *p* and *q* are the probabilities of a T-to-C substitution at each nucleotide for new and old transcripts, respectively, and *n*_*i*_ is the number of uridine nucleotides observed in the transcript *i*. A consensus sequence for each transcript is built by pooling reads with the same UMI barcode and taking the most frequent variant at each position. 10,000 UMIs were randomly sampled and the global substitution probabilities *p* and *q* were estimated based on the above mixture model. The model was fit by maximizing the likelihood function using the Nelder-Mead algorithm. The optimization was repeated ten times with random initialization values for *θ*, *p*, and *q* in the range [0,1], keeping the best fit with *θ* ∈ [0,1].

For mouse cortical neuronal culture datasets, besides Ex and Inh1, we respectively combined 4 inhibitory neuronal clusters (Inh 2-4 and Inh-NP) and 3 non-neuronal clusters (Ex-NP1, Ex-NP2 and RG) to obtain enough UMI for global parameters estimation at each time-point. By doing this, Ex, Inh, and 2 combined clusters were subjected to statistical modelling. For mESC data sets, we assumed that *Tet*-TKO will not affect 4sU utilization efficiency and thus combined WT and TKO data sets to estimate unified global parameters, *p* and *q*, for 3 cell states (pluripotent, intermediate and 2C-like). To this end, 20 sets of *p* and *q* (5 time-points X 4 combined clusters) were determined in cortical neuronal data sets and 3 sets of *p* and *q* (pluripotent, intermediate and 2C-like clusters) were estimated for mESC data sets. Once these global parameters were determined, they were used to estimate the fraction of new transcripts.

##### 1. Computing aggregate new transcripts for cell clusters

**For Figs 2C and S3d**, we aggregate all the UMIs belongs to the same cell-type and estimate the fraction of new transcripts *θ* for each gene with more than 100 UMIs in that cell type at each time-point. The likelihood function for the mixture model above was maximized using the Brent algorithm with the constraint *θ* ∈ [10^<=^, 1]. The 95% confidence interval was calculated from the Hessian matrix, and *θ* estimates for genes with a confidence interval greater than 0.2 were thrown out. The new transcript (N) was then estimated:

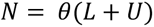

Where *θ* is the fraction of newly-transcribed RNAs for a gene in a cell type, *L* is labeled transcripts of a gene, *U* is unlabeled transcripts of a gene. The pre-existing transcript was calculated by: (1 − *θ*)(*L* + *U*).

##### 2. Computing new transcript for each individual cell

In theory, for every cell the fraction of new transcripts should be calculated for each gene, which requires adequate gene coverage of each individual cells as bulk RNA-seq analysis. However, for data sets generated by high-throughput droplet-based scRNA-Seq methods and shallow sequencing, it is not feasible to obtain sufficient coverage for every gene in individual cells. Moreover, modeling every gene in thousands of cells were computationally intensive. To obtain the single-cell level estimation of new transcripts in each cell, we introduced detection rate *α* to estimate the new transcript for each cell. Since 4sU incorporation is random but each cell may vary in 4sU incorporation and most genes have the highly similar detection rate *α* (**Supplementary Fig. 3d**), we used the strategy above to estimate the fraction of new transcripts for individual cell. The detection rate *α* was then computed by diving all the labeled transcripts of a cell by estimated new transcript of that cell. We discarded the cells without-range values (*α* > 1). We computed the detection rate *α* for 88.3% (18,133/20,547) of mouse cortical cells, whereas 95.1% (6,609/6,952) of mESC cells were retained. The mean of detection rates *α* were 60% and 66% in cortex and mESC, respectively. For each individual gene, the new transcript was computed according to the formula:

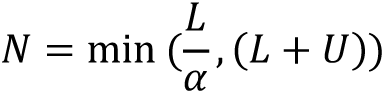

Where *α* is the T-to-C detection rate of a cell, *L* is the number of labeled transcripts of a gene in that cell, *U* is the number of unlabeled transcripts of a gene. The number of pre-existing transcripts was calculated by: *L* + *U* − *N*. The computed new and old transcripts were used for all downstream single-cell-based analysis, including stacked UMAP (**right panel of Fig. 2b** and **Supplementary Fig. 4a**), SCENIC (**Figs. 2d, 3c, 4f, Supplementary Fig. 5** and **Supplementary Fig. 8a**) and new RNA velocity analysis (**Figs. 3a** and **5b**).

#### Gene ontology (GO) enrichment analysis

GO enrichment analysis was performed as previously described^12^. To identify functional categories associated with defined gene lists, the GO annotations were downloaded from the Ensembl database. An enrichment analysis was performed via a hypergeometric test. The *P* value was calculated using the following formula:

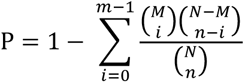

where N is the total number of background genes, n is the total number of selected genes, M is the number of genes annotated to a certain GO term, and i is the number of selected genes annotated to a certain GO term. *P* value was corrected by function *p.adjust* with false discovery rate (FDR) correction in R. GO terms with FDR<0.05 were considered enriched. All statistical calculations were performed in R.

For enrichment analysis of stable/unstable mRNAs (**Supplementary Fig.10c**), genes were ranked by the RNA half-life. Top 10% genes with longest half-lives were defined as stable genes, whereas top 10% genes with shortest half-life were considered as unstable. Then the stable and unstable gene sets were subjected into GO enrichment analysis. For **Fig. 4d**, genes showed >1.5-fold changes between pluripotent and 2C-like state were selected and subjected into GO enrichment analysis.

#### Identification of differentially expressed genes (DEGs)

Differential gene expression analysis of new transcripts between different time-points of neuronal activation (15, 30, 60 and 120 min) and control (0 min) was performed using the function *FindMarkers* in Seurat, using a Wilcoxon rank sum test. New transcripts with a fold-change of more than 1.5 and an adjust *P*-value less than 0.05/4 were considered to be differentially expressed (**Supplementary Fig. 4b**). Neuronal induction genes were defined if a new transcript was significantly increased in at least one time-point with KCl stimulation in any one of cell-types (**Fig. 2c and Supplementary Fig. 4b**). For comparison among 3 cell states within WT mESCs (**Fig. 4e**), genes encoding DNA binding proteins with a fold-change of new transcript expression level more than 0.25 (log-scale) and an adjust *P*-value less than 0.2 were listed.

#### Estimation of RNA half-life

For each gene, we separately aggerate labeled and unlabeled UMI counts in each cell state (**Fig. 6** and **Supplementary Fig.11**). Then the fraction of labeled transcripts was calculated with summed labeled UMI counts divided by total UMI counts (labeled + unlabeled). The fraction of labeled transcripts was normalized to chase-onset (0 min). R function *nls* was used to perform curve fitting with the parameters setting: “*y ∼ I(a*exp(-b*x)+c)*”, “*start=list(a=1, b=1, c=1)*” and “*na.action=na.exclude*”. We kept the fit with the goodness of R^2^ > 0.7.

#### RNA velocity analysis

For standard RNA velocity analysis (splicing RNA velocity), we started with the bam files which were generated by the Drop-Seq computational analysis pipeline. The reads were demultiplexed using dropEst^44^ (version 0.8.5) pipeline, using “-m -V -b -f -L eiEIBA” options to annotate bam files. The genome annotations (mm10, Gencode release vM13) were used to count spliced and unspliced molecules for each experiment. The python package scVelo^21, 45, 46^ (https://github.com/theislab/scvelo, version 0.1.19) were employed to perform RNA velocity analysis. Default parameter settings were used, unless stated otherwise. After loading spliced and unspliced molecules to scVelo, for analysis of excitatory neuron data sets **(left panel of Fig 3a**), genes with less than 5 counts of spliced or unspliced molecules were filtered out. Spliced and unspliced counts of top 1,000 highly variable genes were KNN-imputed in a PCA reduced space with 10 components, using 15 neighbors and ‘distances’ mode. For new RNA velocity analysis (using the ratio of new over total transcripts), all the parameters are the same as splicing RNA velocity analysis except that we loaded new transcripts (as unspliced counts) and total transcript (as spliced counts) into scVelo (**right panel of Fig 3a**). For new RNA velocity analysis of mESCs (**Fig 5a**), top 3,000 highly variable genes were KNN-imputed in with 10 PCs, using 15 neighbors and ‘distances’ mode as well. The gene-specific velocities are obtained by fitting a ratio between new and total mRNA by function ‘*scv.tl.velocity*’. Finally, function ‘*scv.pl.velocity_embedding_stream*’ was used to project the velocities onto UMAP.

To directly compare splicing and new RNA velocity, we asked which of them could best predict whether well-known activity induced genes upregulated with different durations of KCl stimulation in excitatory neurons. The list of 137 primary response genes was previously defined by bulk RNA-Seq^16^. We calculated the average induction fold of genes (expressed in >1% cells) at 30 min of KCl stimulation compared to 0 min (**Fig 3b**). For splicing RNA velocity, we used unspliced counts to calculate the induction fold while estimated new transcript were used in new RNA velocity analysis. Finally, *auc* function in R package pROC was conducted to calculate the area under receiver operating characteristic curve (AUC), which reflects the accuracy of predicting the known PRGs by gene induction fold computed in different method. The plots were generated by function *plot.roc* in pROC package^47^.

#### Gene regulatory network (GRN) analysis by SCENIC

To assess the regulatory activity of transcript factors associated with different cell states or cell-types, we used SCENIC^18^ (version 1.1.2.2) to perform gene regulatory network (GRN or regulon) analysis.

Regulatory modules are identified by inferring co-expression between TFs and genes containing TF binding motif in their promoters. We separate expression matrix into two parts based on the expression level of new and old transcripts, then combined them as inputs to SCENIC analysis, which enabled us to identify specific regulatory modules associated with either new or old transcriptomes from the same cell. Two gene-motif rankings, 10kb around the TSS and 500 bp upstream, were loaded from RcisTarget databases (mm9). Gene detected in > 1% of all the cells and listed in the gene-motif ranking databases were retained. To this end, 8,744 genes in cortical culture data and 9,388 mESC genes were subjected into the downstream analysis. Then GRNBoost, which was implemented in pySCENIC, was used to infer the co-expression modules and quantify the weight between TFs and target genes. Targets genes that did not show a positive correlation (> 0.03) in each TF-module were filtered out. SCENIC found 4,944 and 5,406 TF-modules in mouse cortical neuronal culture and mESC data sets, respectively. A *cis*-regulatory motif analysis on each of the TF-modules with RcisTarget revealed 277 and 325 regulons in cortical culture and mESC data, respectively. The top 1 percentile of the number of detected genes per cell was used to calculate the enrichment of each regulon in each cell. For **Figs. 2d, 4f, and Supplementary Fig. 8**, we computed the mean AUC of all cells belongs to defined groups, then scaled the mean AUC by function *scale* in R. R package pheatmap^48^ was used to draw the heatmap.

For **Fig. 4f and Supplementary Fig. 8**, AUC values of TFs were obtained and then subjected to Wilcoxon rank sum test to access significance of the difference of TF activity. TFs with a fold-change of mean AUC values more than 1.5 and an adjust *P*-value (Bonferroni corrected) less than 0.05 were considered differentially regulated.

#### RNA binding protein motif enrichment analysis

For **Fig. 6f**, 3’ UTR sequences of 552 genes were retrieved from Ensembl Biomart^49^. All the 3’ UTR sequences of a gene were obtained to explore all possible binding site. The position weight matrix (PWM) of 188 mouse RNA binding protein motifs were downloaded from CISBP-RNA database^50^ (http://cisbp-rna.ccbr.utoronto.ca/). FIMO^51^ (version 5.0.5) motif scanning software in MEME Suite was used to search for the RNA binding motifs in the 3’ UTR sequences. 3’ UTR sequences with *P*-values smaller than the default threshold (0.0001) were considered to contain a binding motif. As a result, 533 genes were identified to have RNA binding motif in their 3’ UTR. Hypergeometric test was performed in R by function *phyper*, which was used to access significance of RNA binding motif enrichment in defined gene sets.

The pairs of RNA binding protein and targets were visualized in Cytoscape.

## Data availability

All sequencing data associated with this study will be available through the Gene Expression Omnibus (GEO) database.

## Code availability

The analysis source code underlying the final version of the paper will be available on GitHub repository (https://gibhub.com/wulabupenn/scNT-seq).

---

**Supplementary Figure 1:**
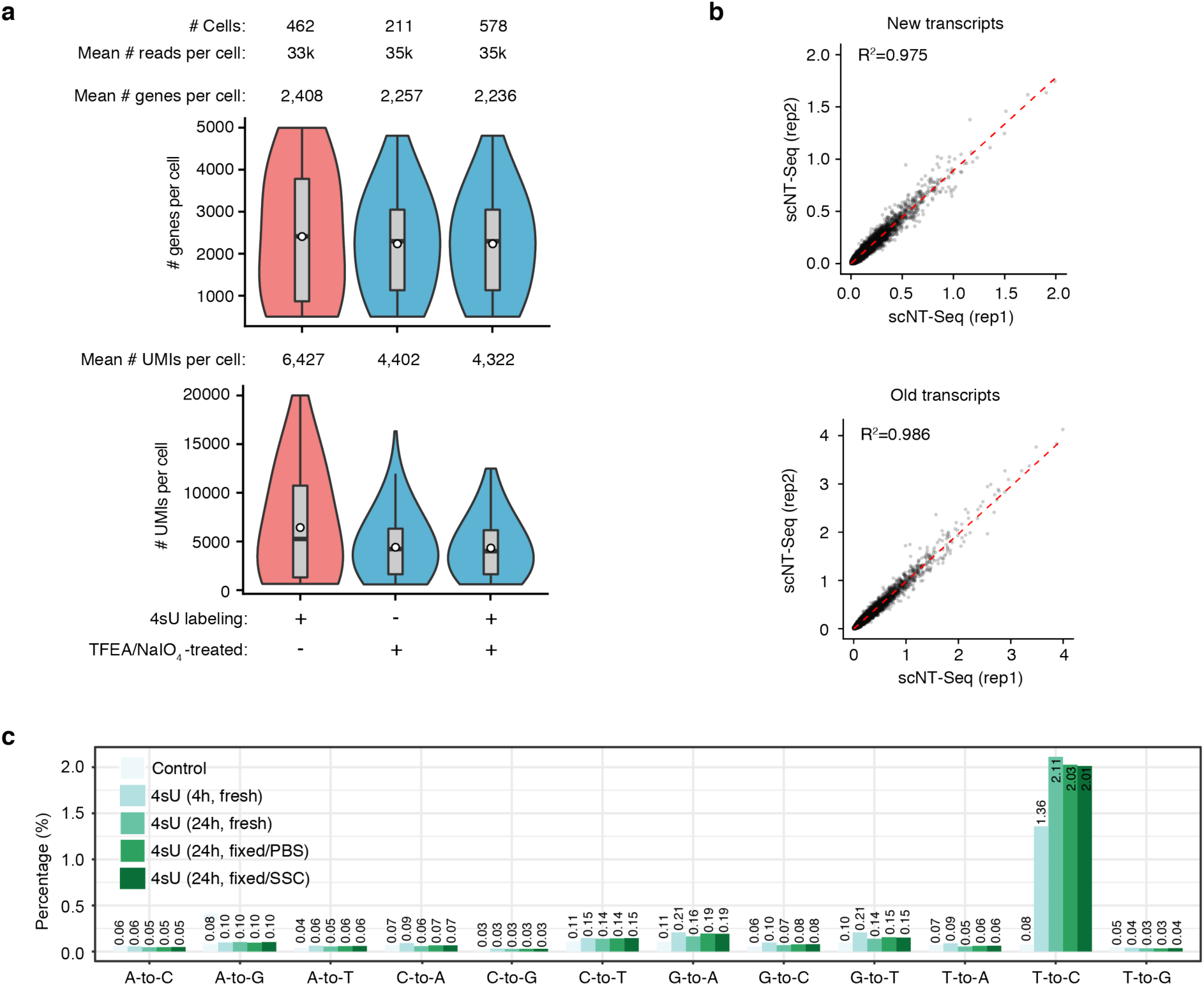
Quality control of scNT-Seq in mESCs. **a.** Violin plot comparing the number of genes and number of UMIs detected in individual cells among different treatment conditions. Cell number and sequencing depth were shown on the top. **b.** Scatter plot showing reproducibility between two replicates. **c.** Transcriptome-wide nucleotide substitution rates reveal the effect of labeling time (100 μM 4sU/4hr or 200 μM 4sU/24hr) and methanol fixation (two rehydration buffers, PBS versus SSC) on T-to-C conversion rate. The control sample (100 μM 4sU/4hr) was not treated with TFEA/NaIO_4_.

**Supplementary Figure 2:**
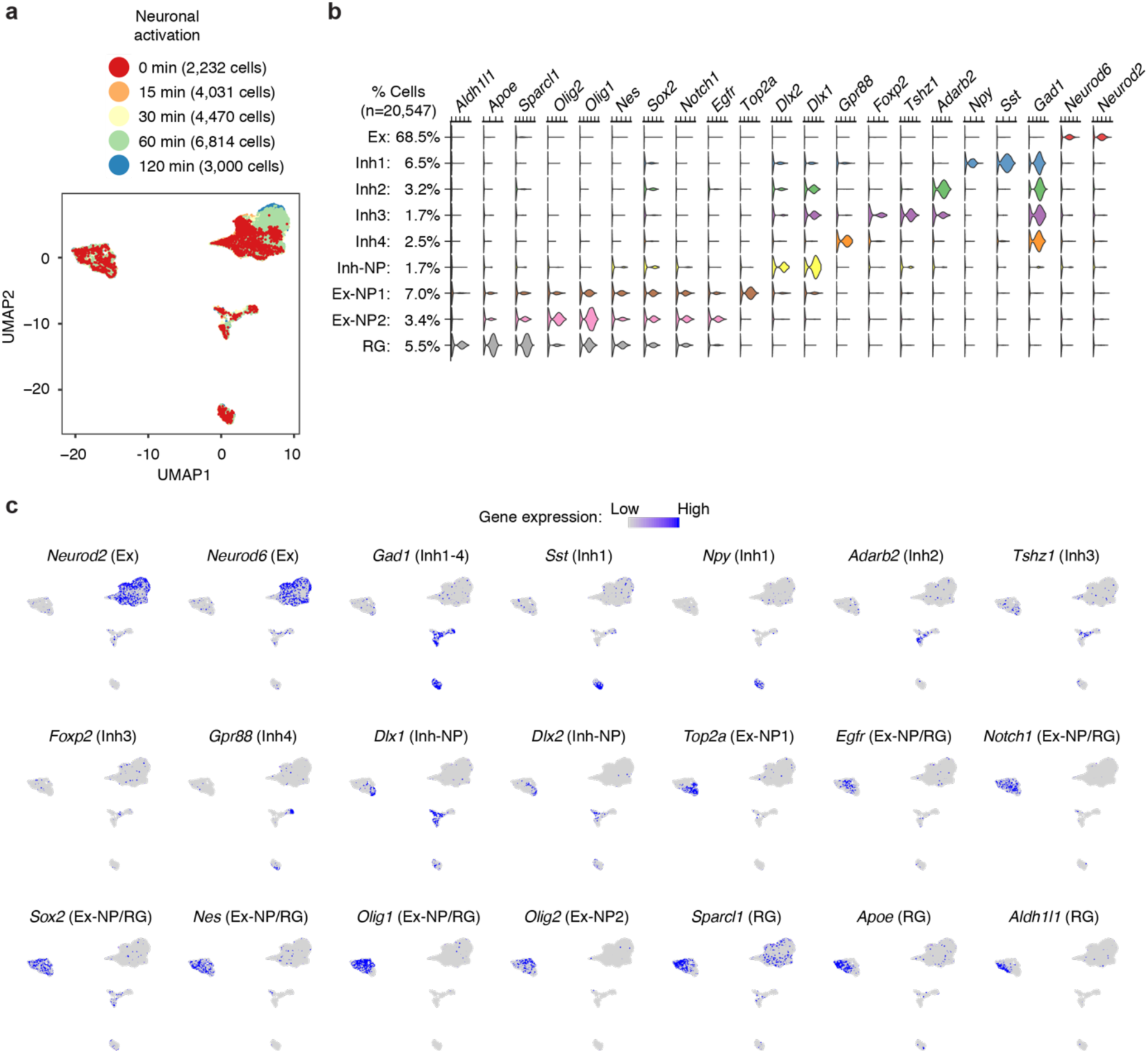
scNT-Seq identifies different cell types in primary mouse cortical cultures. **a.** UMAP visualization of 20,547 cells from mouse cortical cultures (same UMAP plot as Fig. 2b). The cells are colored by different durations of neuronal activation. **b.** Violin-plot showing gene expression levels of representative marker genes in different cell-types. **c.** Marker gene expression is scaled by colors in the same UMAP as (a). Ex, excitatory neurons; Inh, inhibitory neurons; NP, neural progenitors; RG, radial glial progenitor cells.

**Supplementary Figure 3:**
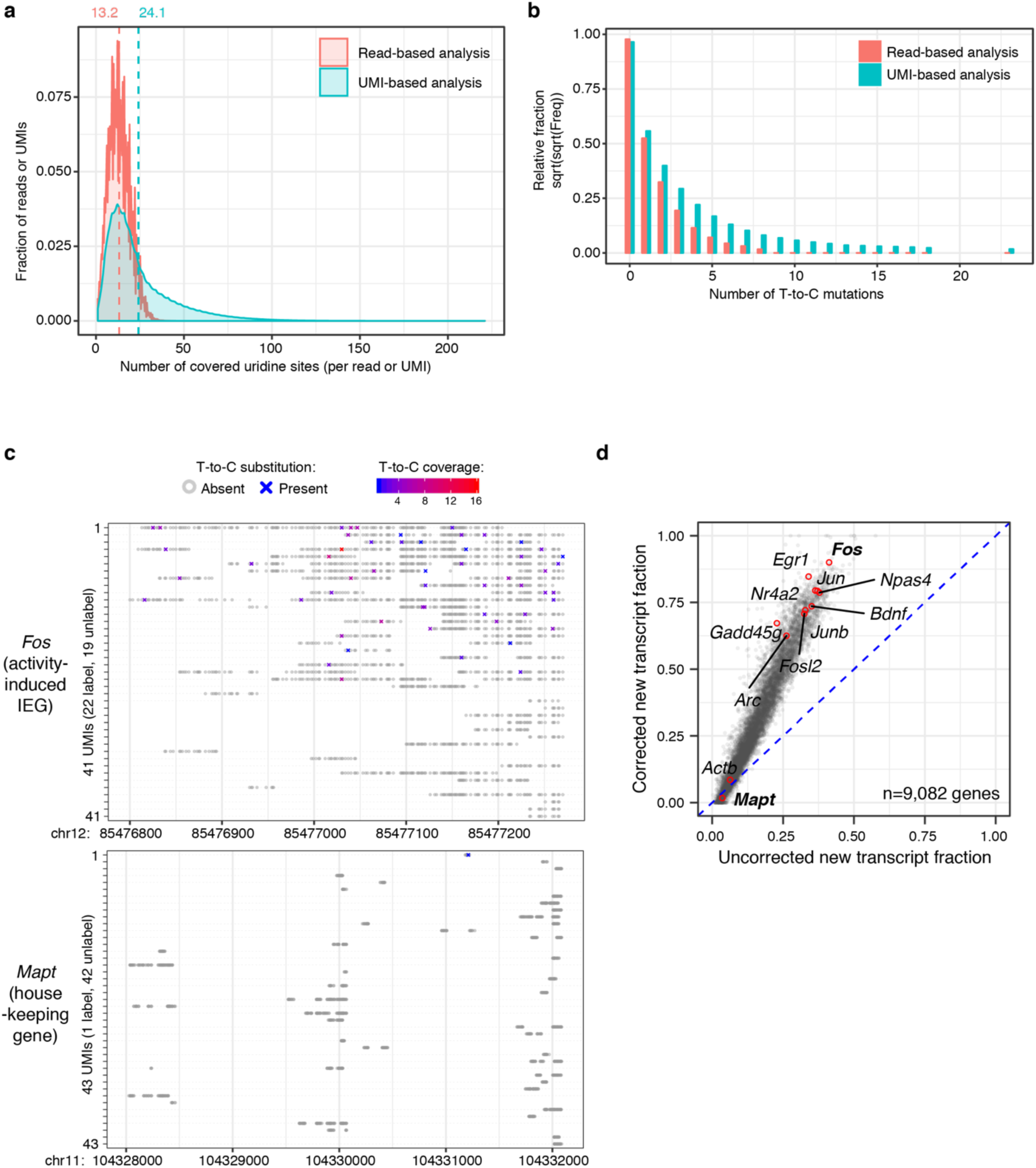
UMI-based statistical correction of fractions of newly-transcribed RNAs. **a.** Density plot of the number of covered uridine sites per read (60 bp) or UMI. Data set of excitatory neurons with 60 min KCl stimulation were shown. **b.** Bar plot of the number of T-to-C substitutions per read (60 bp) or UMI. Data set of excitatory neurons with 60 min KCl stimulation were shown. **c.** All UMIs of the *Fos* and *Mapt* genes from one excitatory neuron with 60 min KCl stimulation were shown. Grey circles stand for T without T-to-C conversion, while crosses (“X”s) denote sites of T-to-C conversion in at least one read. **d.** Comparison of new RNA levels of each gene in excitatory neurons (with 60 min KCl stimulation) before and after statistical correction. Ten neuron activity-dependent genes and two house-keeping genes are highlighted with red dots.

**Supplementary Figure 4:**
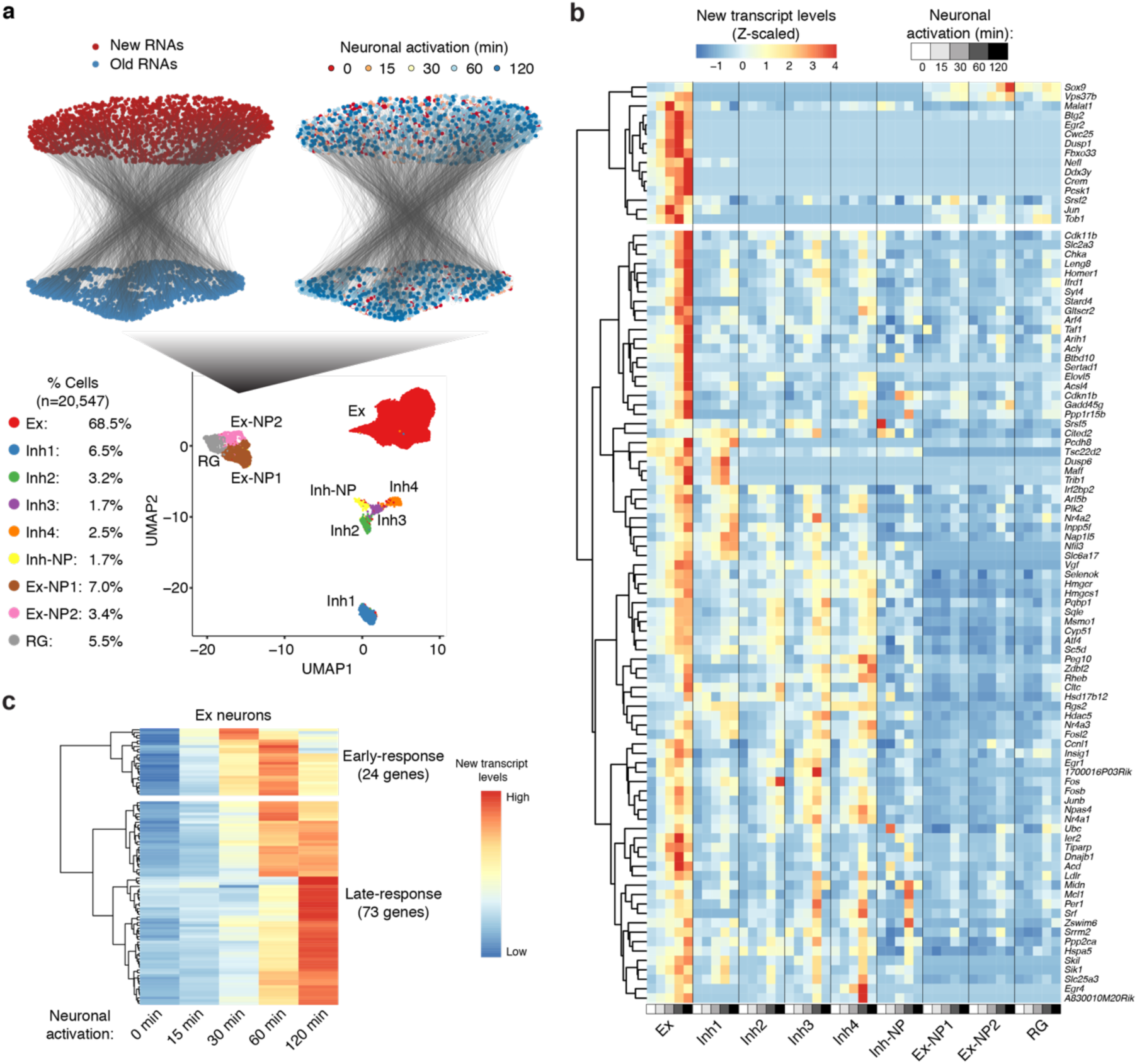
Different cell types show different neuron activity-dependent gene expression programs. **a.** UMAP visualization of the expression profiles of 20,547 cells from primary mouse cortical cultures with or without KCl stimulation (bottom panel, same UMAP plot as Figure 2b). Cells from non-neuronal cell clusters (RG/Ex-NP) are sub-clustered based on newly-transcribed or pre-existing mRNAs (top). The newly-transcribed and pre-existing mRNAs of same cell were connected by black line. Ex, excitatory neurons; Inh, inhibitory neurons; NP, neural progenitors; RG, radial glial progenitor cells. **b.** Heat map showing new transcript levels of genes with significantly increased expression level after KCl stimulation in at least one cell type. **c.** Heat map showing new transcript levels of early- and late-response genes in excitatory neurons with different durations of KCl stimulation. 97 significantly induced genes were clustered into 2 groups (early- and late-response).

**Supplementary Figure 5:**
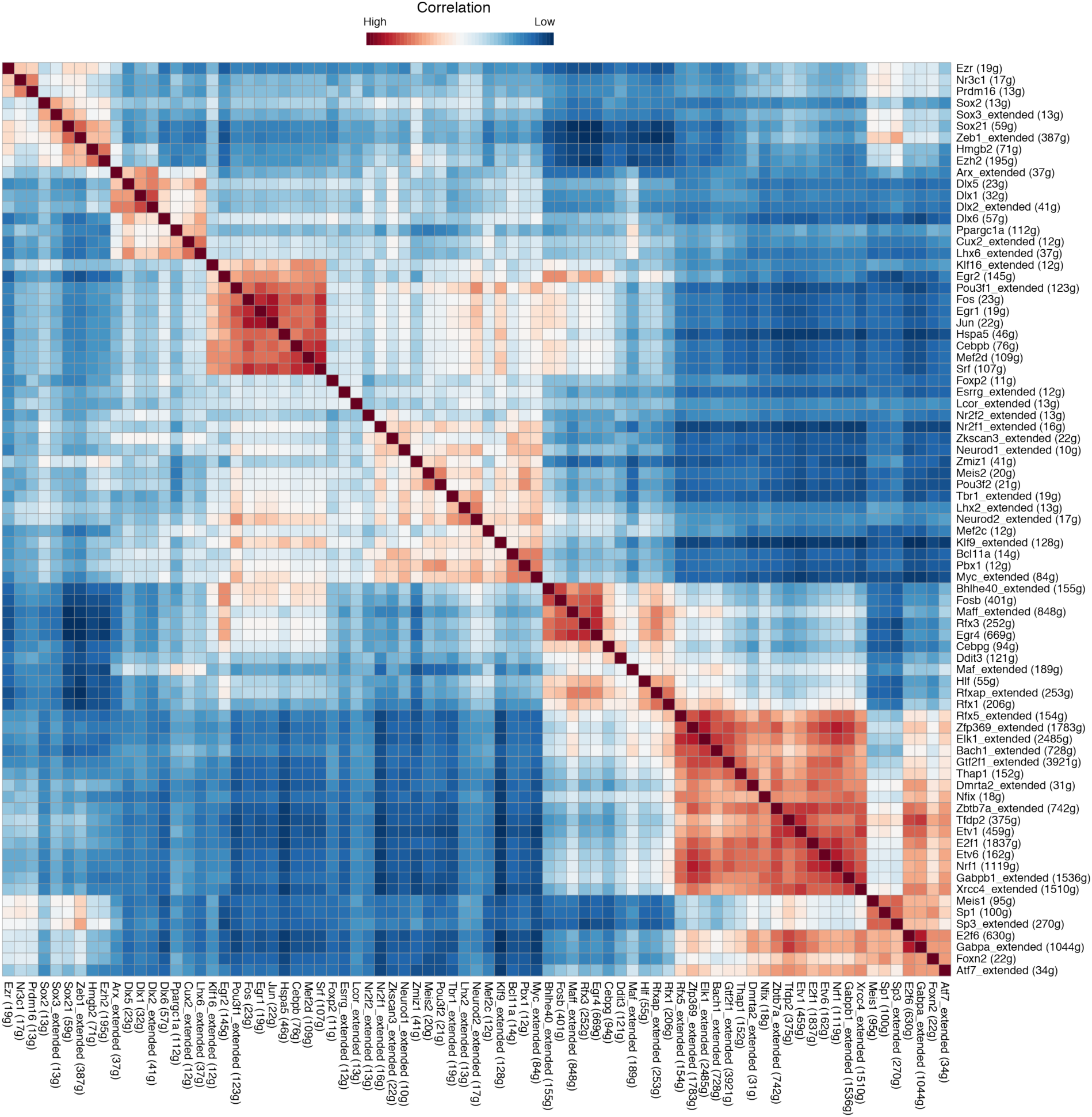
Heat map of activity-regulated TF regulons in mouse cortical cultures. Heat map showing correlated transcription factor activity (quantified by SCENIC) during neuronal activity-dependent gene expression.

**Supplementary Figure 6:**
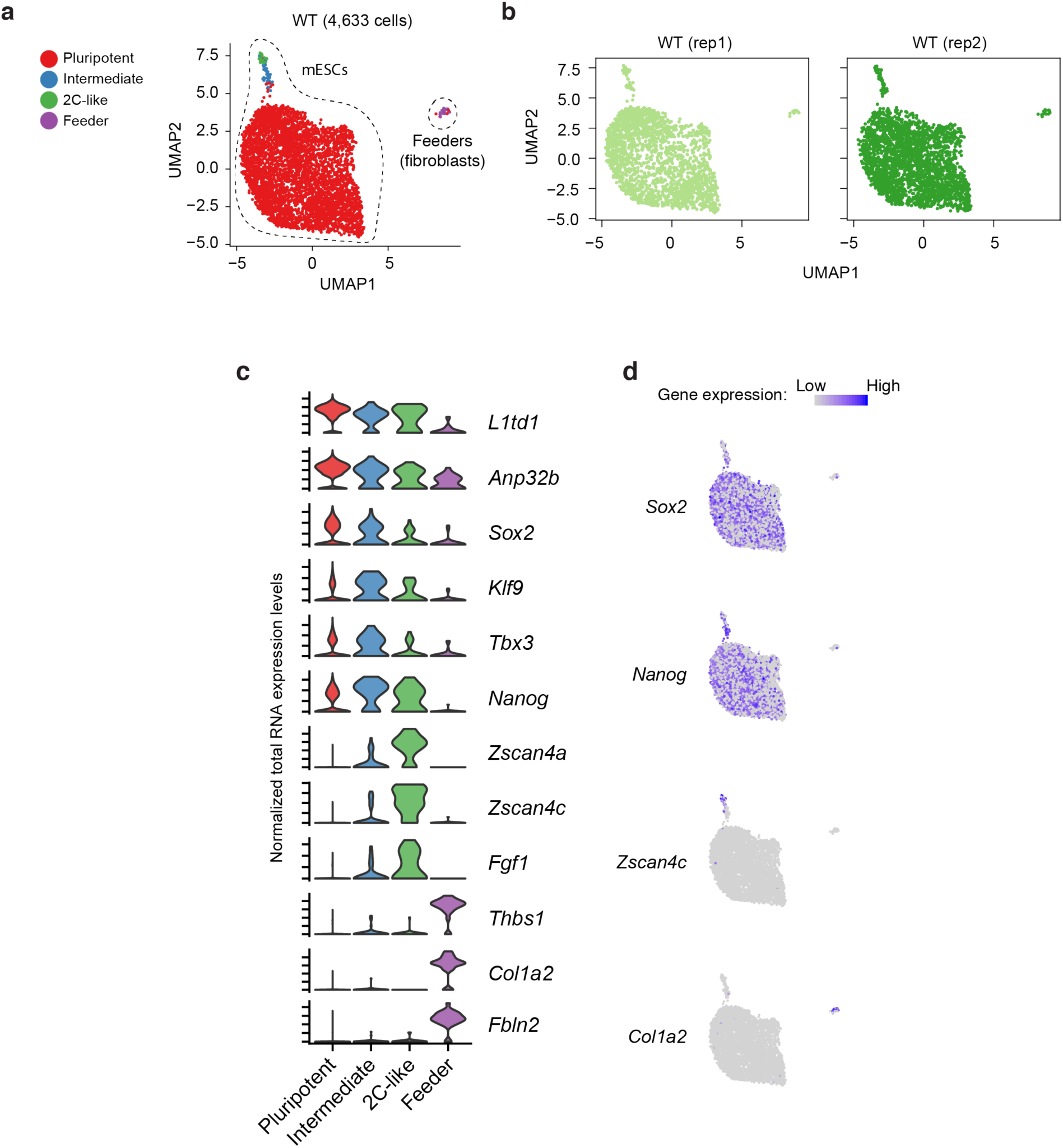
scNT-Seq reveals different stem cell states in mESC cultures. **a.** UMAP visualization of 4,633 WT mESCs from two replicates. The cells are colored by different cell-types or cell-states. Feeders (purple dots) are contaminated mouse embryonic fibroblasts in cell culture. **b.** UMAP plots were colored by replicate. **c.** Violin plot showing gene expression level of marker genes in different stem cell states or cell-types. **d.** Feature-plots showing marker gene expression is scaled by colors in the same UMAP as (a).

**Supplementary Figure 7:**
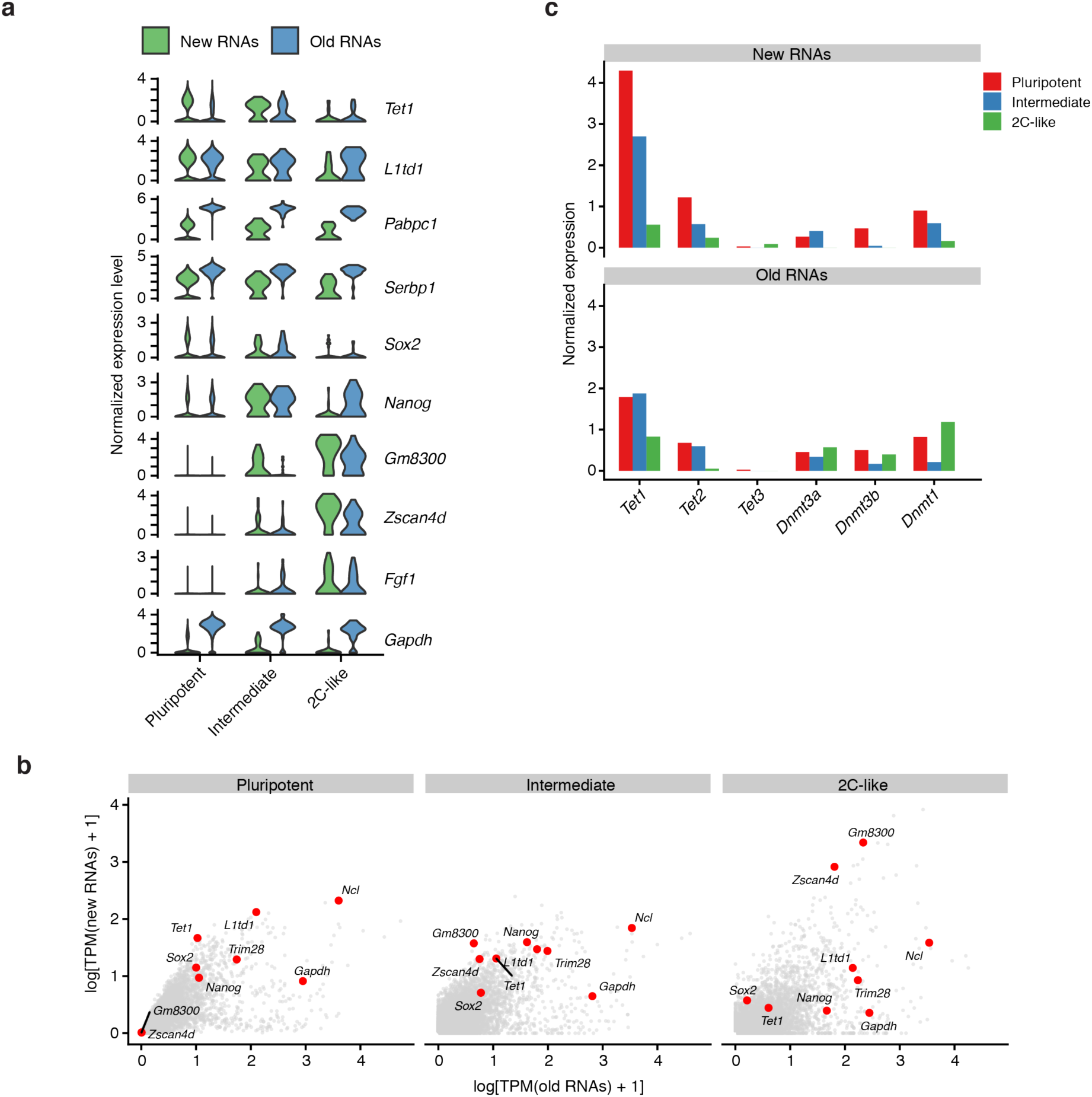
scNT-Seq reveals newly-transcribed and pre-existing RNA level of representative genes in mESCs. **a.** Violin plots showing expression levels of newly-transcribed and pre-existing transcripts of selected genes in three cell-states of mESCs. **b.** Scatterplots showing newly-transcribed and pre-existing RNA levels of representative genes, including pluripotent genes (*Sox2, Nanog*), 2C-like state specific genes (*Zscan4d, Gm8300*), and house-keeping gene (*Gapdh*). **c.** Newly-transcribed and pre-existing RNA levels of major DNA methylation regulators in three stem cell states.

**Supplementary Figure 8:**
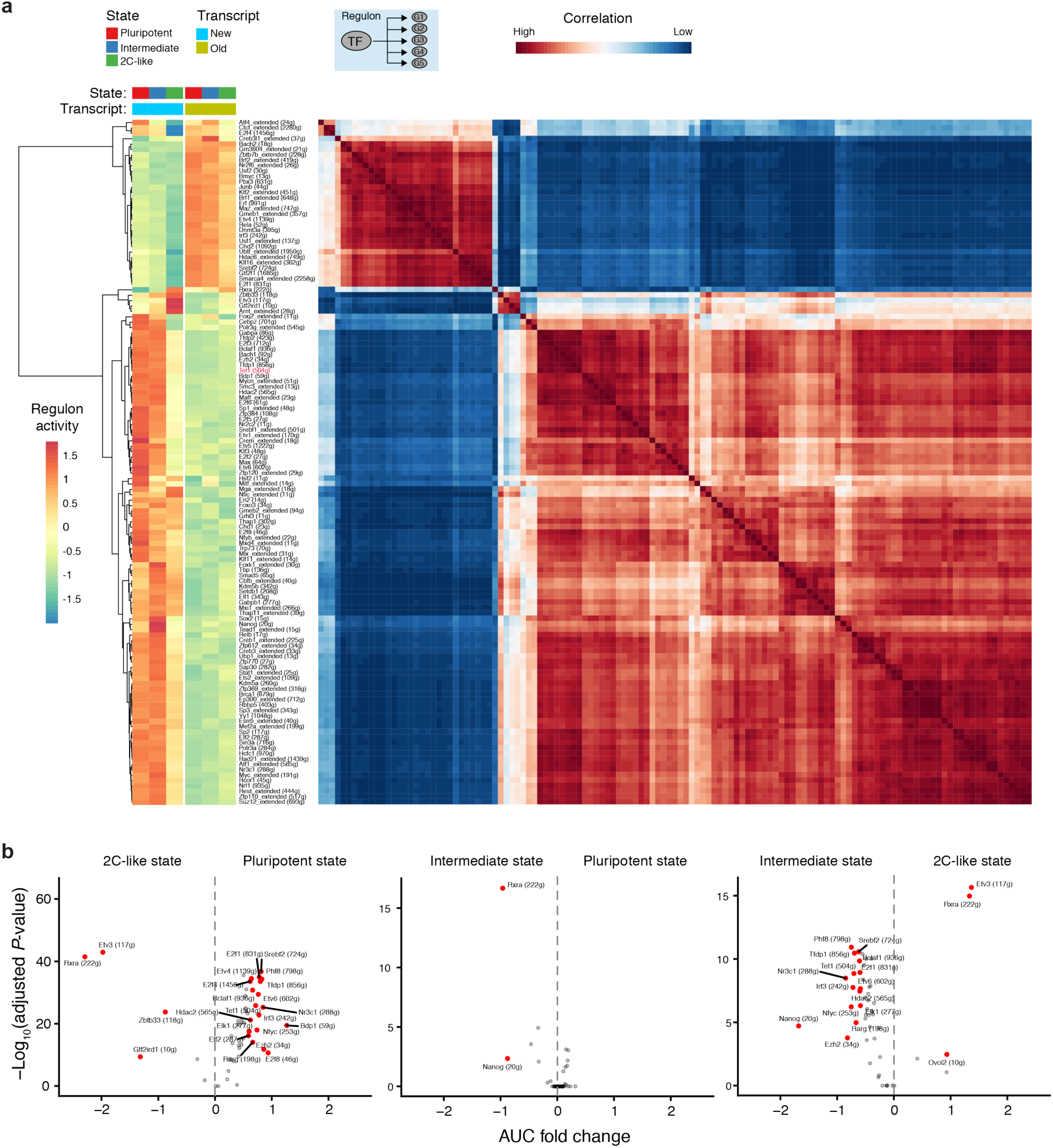
scNT-Seq reveals TF activity changes during pluripotent-to-2C transition in mESCs. **a.** Heat maps showing TF regulon activity (left) and the similarity of transcription factors activities in three stem cell states (right). **b.** Volcano-plot showing state-specific TF regulon activity. TFs with a fold-change of mean AUC values more than 1.5 and adjust *P*-value less than 0.05 were highlighted in red.

**Supplementary Figure 9:**
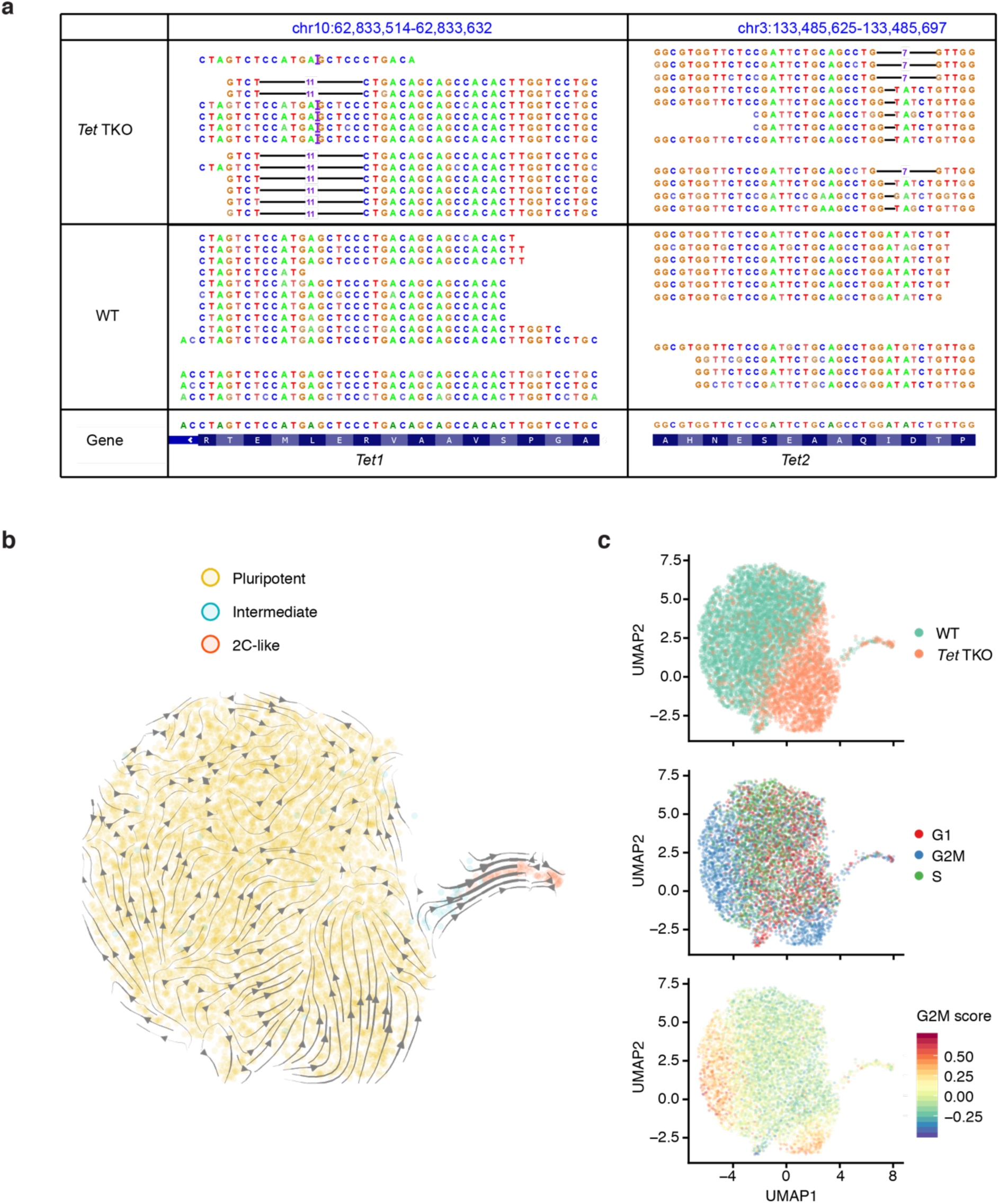
scNT-Seq analysis of TET-dependent regulation of the plutipotent-to-2C transition in mESCs. a. Validation of genotypes of the *Tet1* (-11bp/+1bp) and *Tet2* (-7bp/-1bp) genes in *Tet*-TKO cells by aligning scNT-Seq reads to the CRISPR/Cas9 genome editing sites. b. Projection of cell states onto the same UMAP plot as Fig. 5a. c. Projection of genotypes and the cell-cycle states onto the same UMAP plot as Fig. 5a.

**Supplementary Figure 10:**
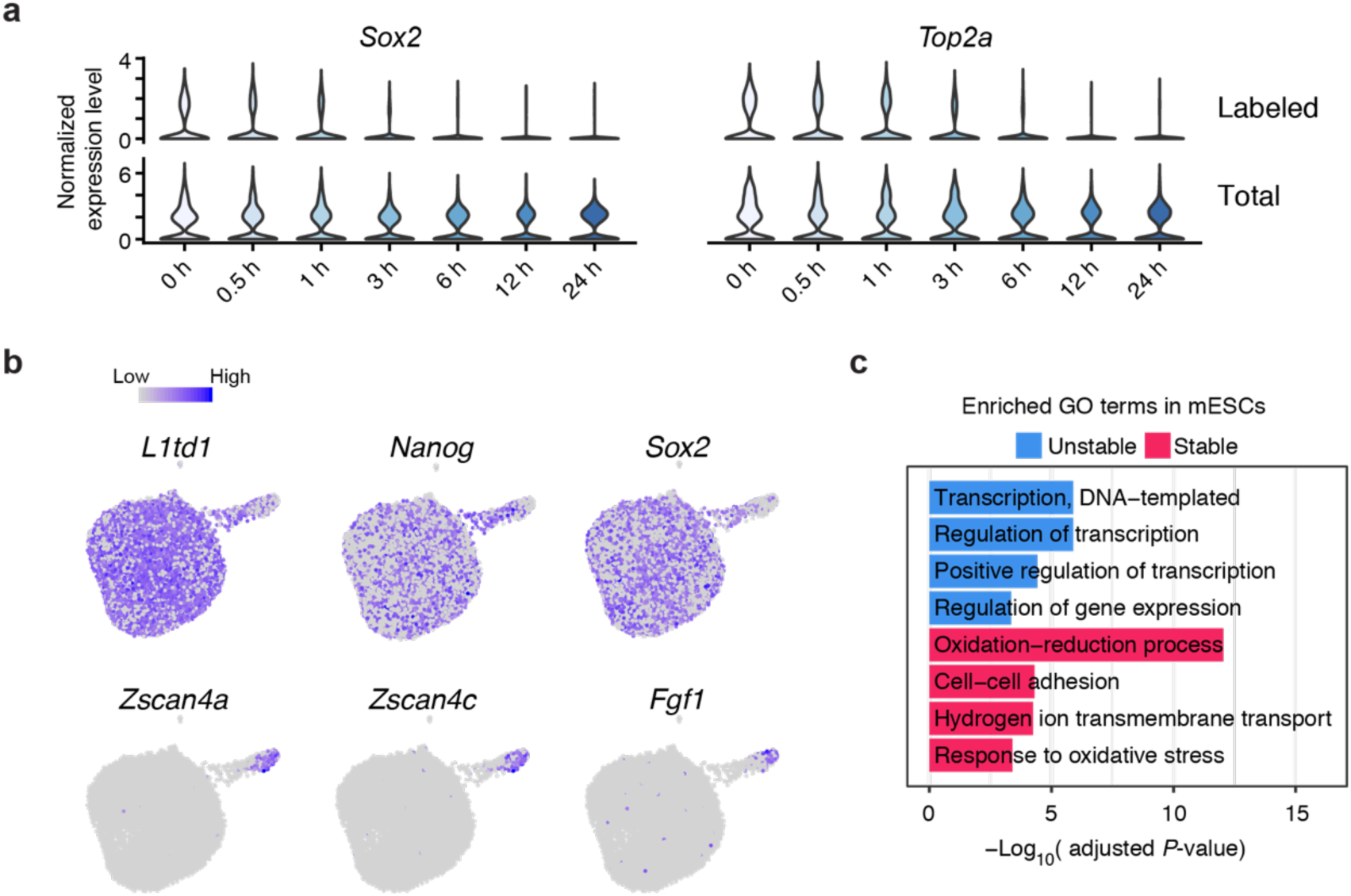
Pulse-chase scNT-Seq reveals transcript-specific mRNA decay of mESCs. **a.** Violin plots showing levels of labeled and total transcripts of the *Sox2* and *Top2a* genes during pulse-chase assay. **b.** Marker gene expression is scaled by colors in the same UMAP plot as Fig. 6b. **c.** Enrichment analysis of GO terms within stable (top 10% genes with longest half-lives) and unstable genes (top 10% genes with shortest half-life) in pluripotent state mESCs.

**Supplementary Figure 11:**
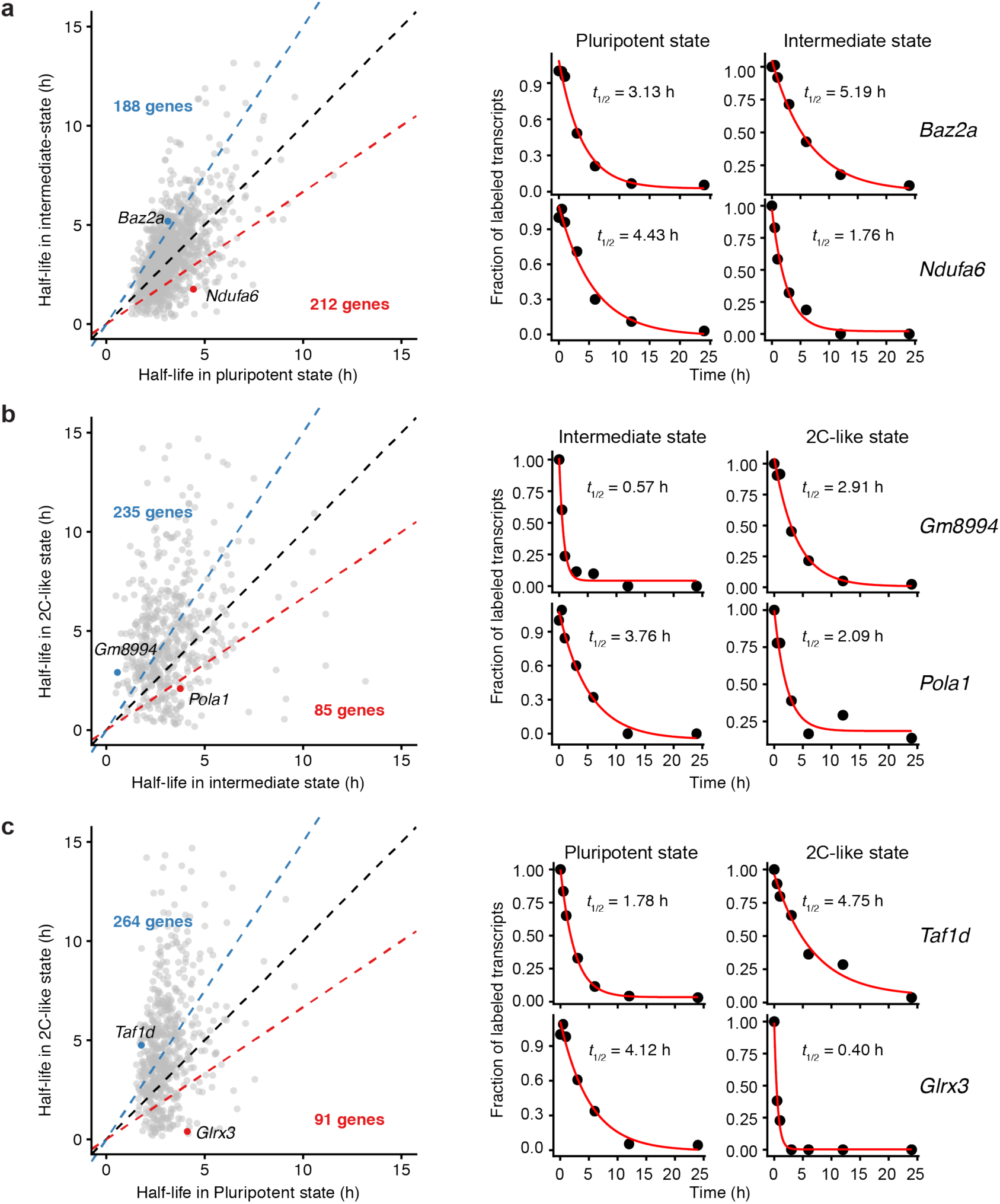
Pulse-chase scNT-Seq reveals cell state-specific mRNA decay in mESCs. Scatterplots comparing the RNA half-life of commonly detected transcripts between two stem cell states (**a**: pluripotent vs. intermediate; **b**: intermediate vs. 2C; **c**: pluripotent vs. 2C) and number of genes showing >1.5-fold change in RNA half-life (indicated by blue and red dashed line) between two states are shown (left panels). Representative genes showing state-specific RNA half-life are shown in the right panels.

**Supplementary Figure 12:**
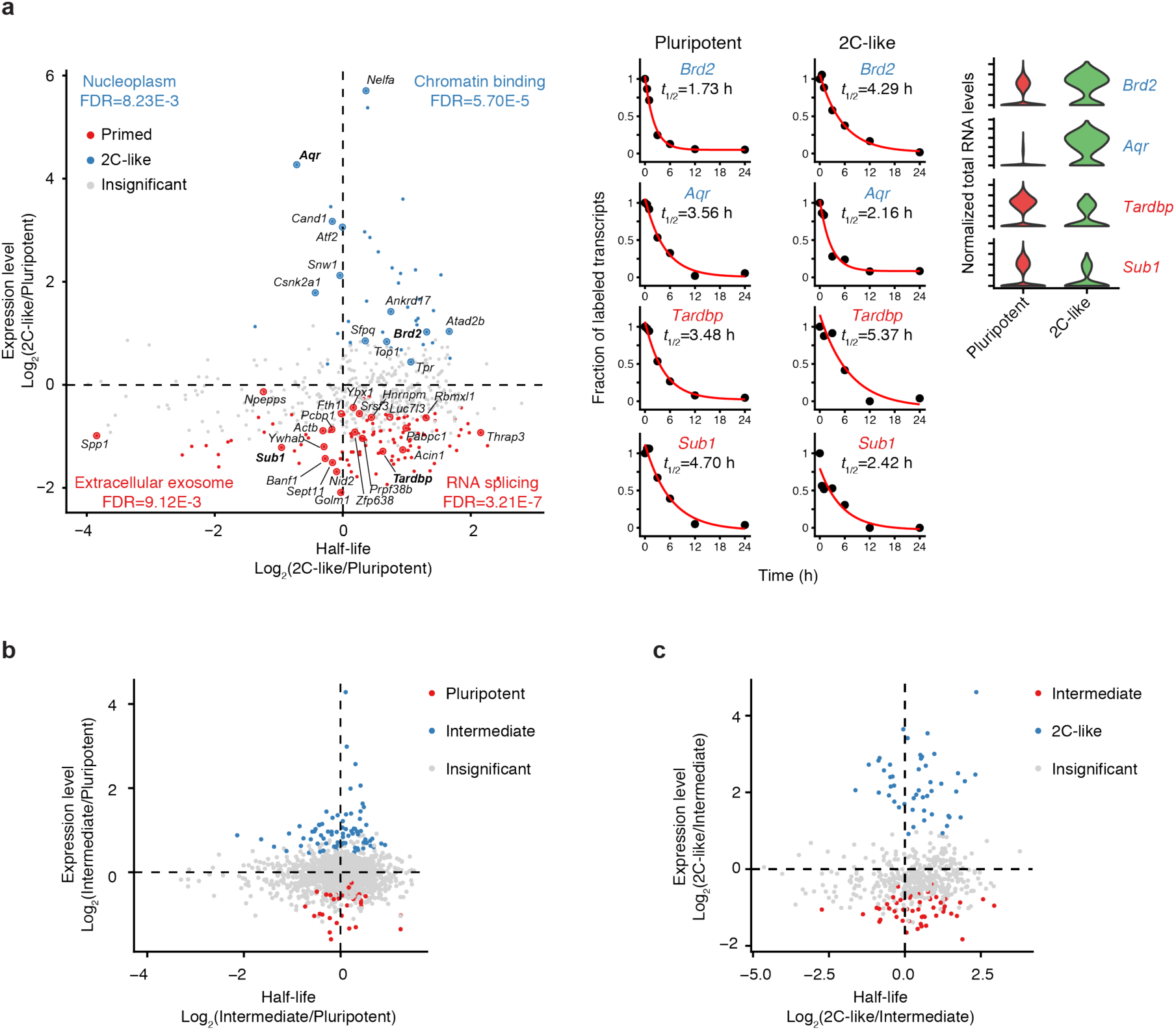
Characterization of mRNA stability and gene expression levels in different states of mESCs. **a.** Scatterplot showing the correlation between gene expression levels (total transcripts) and RNA half-life in pluripotent and 2C-like states. Representative GO terms enriched in genes in each quadrant, as well as genes belong to these GO terms, are highlighted. The mRNA decay kinetics and expression level of selected 4 genes were also shown in the middle panel. Representative state-specific genes with different RNA half-life are shown in the right panel. 2C-like state enriched genes are highlighted in blue, while genes preferentially expressed in the pluripotent state are highlighted in red. **b-c.** Scatterplot showing the correlation between gene expression levels (total transcripts) and RNA half-life in different states (b: Pluripotent vs. intermediate; c: intermediate vs. 2C-like). Genes that are differentially expressed in one stem cell state are highlighted in red or blue.

